# The Origin, Diagnosis, and Prognosis of Oligomannose-Type Diffuse Large B-Cell Lymphoma

**DOI:** 10.1101/2025.03.11.642396

**Authors:** Dylan J. Tatterton, Maddy L. Newby, Joel D. Allen, Benjamin Sale, Giorgia Chiodin, Patrick J. Duriez, John Butler, Katy J. McCann, David W. Scott, Ryan D. Morin, Kostiantyn Dreval, Andrew J. Davies, Dean J. Bryant, Max Crispin, Francesco Forconi

## Abstract

The acquisition of N-glycosylation sites occupied by oligomannose-type glycans in the immunoglobulin complementarity-determining region (CDR) is an early clonal tumor-specific identifier of follicular lymphoma (FL). CDR-located N-glycosylation sites are also acquired in germinal-center-B-cell-like diffuse large B-cell lymphomas (GCB-DLBCL), but their significance is less defined. We used RNA-seq immunoglobulin assembly to determine the frequency and CDR location of the acquired N-glycosylation sites (AGS) in two large independent DLBCL cohorts. Composition of the glycans occupying the AGS was determined using liquid chromatography-mass spectrometry and correlated with cell-of-origin, FL signature (defined by EZB phenotype or *BCL2* translocation), transcript profile, and clinical outcome. CDR-located AGS were observed in 41-46% of GCB-DLBCL but were rare in other DLBCL. Only CDR-located AGS of DLBCL with an FL signature were occupied by oligomannose-type glycans. These DLBCL were termed Mann-type DLBCL. Conversely, the AGS of the other DLBCL were either non-glycosylated or occupied by complex-type glycans. Mann-type status was an independent marker of short progression-free survival and overall survival. In contrast, the other GCB-DLBCL cases, including those with an FL signature but without AGS, had the best outcomes. Mann-type DLBCL overexpressed gene-sets of cell growth, survival, and cycling, and underexpressed proinflammatory and apoptotic pathways, irrespective of concomitant *MYC* translocations. Acquisition of Mann-type glycans is a highly selective environmental pressure, identifying the most aggressive GCB-DLBCL with an origin related to FL. The detection of AGS in the CDR of GCB-DLBCLs with a BCL2 translocation defines Mann-type DLBCLs, refines prognosis and marks a precise tumor interaction to block early therapeutically.

**Key points:**

- Acquisition of Mann-type glycans is a tumor-specific environmental pressure identifying the most aggressive GCB-DLBCL with an origin related to FL.
- Mann-type DLBCL can be diagnosed by the concomitant detection of AGS in the CDR and a BCL2 translocation.

## Introduction

The B-cell receptor (BCR) is essential for the survival of most mature normal and cancer B cells, and its structure masters their interactions with the microenvironment and fate.^1–3^ Understanding of the BCR structure and function in chronic lymphocytic leukemia (CLL) has led to the development of BCR-associated kinase inhibitors and divided CLL into 2 types with unmutated BCR and bad prognosis or mutated BCR and good prognosis.^4^

In follicular lymphoma (FL), the BCR undergoes a specific transformation from recognizing antigens to engaging with local microenvironmental lectins, promoting the survival and proliferation of FL cells.^5^ This transformation is driven by the presence of oligomannose-type (Mann-type) glycans in the BCR’s complementary-determining regions (CDR), which prevent the maturation of these glycans, likely due to steric hindrance, maintaining their immature state.^6^ These Mann-type glycans interact with the C-type lectin DC-SIGN,^7,8^ found on interfollicular macrophages and follicular dendritic cells of FL.^5,9–11^ This antigen-independent interaction triggers continuous low-level signaling that aids in tissue retention, protects lymphoma cells from apoptosis,^12,13^ and leads to the upregulation of MYC protein and transcript.^8,14^

In diffuse large B-cell lymphomas (DLBCL) the BCR is typically expressed at high levels, but the influences on sIg signaling may be distinct among individual subsets. Gene expression profiling has identified three main subsets based on putative cell-of-origin: germinal center B-cell-like DLBCL (GCB-DLBCL), in which the tumor cells resemble normal germinal center B cells, activated B-cell-like DLBCL (ABC-DLBCL), resembling B cells activated by anti-IgM *in vitro*, and an otherwise “unclassifiable” subset.^15,16^ Genomic studies have further subdivided DLBCL into seven Lymphgen subtypes.^17^ This subdivision seems important for ABC-DLBCL, of which 2 subtypes, MCD and N1, have shown sensitivity to BCR-associated kinase inhibitors.^18^ The most common subtype in GCB-DLBCL is EZB, which is defined by *a* translocation of the *BCL2* gene in chromosome 18 to the *IG* gene locus in chromosome 14, and by mutations in epigenetic regulator genes, including *EZH2* and *CREBBP*.^17,19^ While the mutations of epigenetic regulators can be found in different subtypes, the *BCL2* translocation is exclusive to EZB. The *BCL2* translocation is also the founding hallmark of FL,^20–22^ suggesting a relationship between FL and EZB.^17^ In DLBCL, it also associate with a particularly poor prognosis when a *MYC* translocation coexists.^17,23–25^

However, DLBCL B cells also require to interact with environmental elements at tissue sites, where they typically develop and reside. The influences on BCR signaling in ABC-DLBCL include autoantigens, many interacting with *IGHV4-34*, and mutations increasing the amplitude of BCR signaling.^26,27^ In contrast, ~50% of GCB-DLBCL acquire N-glycosylation sites selectively in the CDR.^6^ However, the consequences and clinical significance of the AGS in GCB-DLBCL are less clear.

In this study, we analyzed DLBCL-derived BCR glycans occupying the acquired N-glycosylation sites (AGS) and investigated the association of glycan composition with location, RNA-seq, genetic, and clinical data of 590 DLBCL from two independent international cohorts using a discovery-validation approach. The data reveal that the occupation of BCR by Mann-type glycans can be predicted by the combination of AGS in the CDR and EZB status (defined by BCL2 translocation), identify a progenitor common with FL, and is an independent marker of poor outcome with the current standard therapies.

## Methods

### Patient cohorts, RNAseq, immunoglobulin gene analysis, and data selection

Two independently validated and publicly available cohorts of DLBCL were used. The BC Cancer Agency (BCCA) de-novo DLBCL cohort was used for discovery (**Table S1A**),^23^ and the National Cancer Institute (NCI) DLBCL cohort was used for validation (**Table S1B**).^28^ Data were obtained from the EGA (BCCA dataset ID: EGAD00001003783, cohort) or the genomic Data commons for Genotypes and Phenotypes (accession phs001444.v1.p1, NCI cohort), and used for RNAseq analysis as described in the **Supplementary methods**. The metadata for the cohorts were obtained as described in the **Supplementary methods**. The full *IGHV-IGHD-IGHJ-IGHC* functional rearranged transcript was obtained from the RNAseq data using our IgSeqR assembly pipeline.^6,29^ Only the cases with full *IG* transcript and clinical data available were used, while the other cases were excluded from the study.

### Production of recombinant lymphoma-derived F(ab)s

Recombinant F(ab)s containing the *IGHV-IGHD-IGHJ* and *IGKV-J/IGLV-J* sequences from FL and DLBCL cases were produced, with sequences from 35 AGS+ve cases from the NCI DLBCL and 12 FL from the University of Southampton (**Table S2**).^30^ The lymphoma-derived F(ab)s were produced using a MEXi HEK-293E cell expression system and were purified using affinity-based chromatography, as described (**Supplementary methods**).^6^

### Glycopeptide mass spectrometry of lymphoma-derived F(ab)s

Separate 50 μg aliquots of purified F(ab)s were digested with trypsin, chymotrypsin (Promega), and/or α-lytic protease (NEB) at a 1:30 (w/w) ratio in 50 mM Tris/HCl, pH 8.0 to generate glycopeptides containing a single N-glycosylation site. Glycopeptides were purified using C18 Zip-tip (Merck Millipore), dried, resuspended in 0.1% formic acid, and analyzed by liquid chromatography-mass spectrometry with an Ultimate U3000 connected to an Orbitrap-Eclipse mass spectrometer (Thermo Fisher Scientific). Peptide and glycopeptide ions were fragmented using stepped higher-energy collisional dissociation fragmentation. Peptides were separated using an Easy-Spray PepMap RSLC C18 column (75 µm x 75 cm) with a 0-32% acetonitrile linear gradient in 0.1% formic acid for 240 minutes followed by 80% acetonitrile in 0.1% formic acid for 35 minutes (flow rate 200 nl/minute; spray voltage 2.5 kV; capillary temperature 275°C). Glycopeptide fragmentation data were extracted using Byos v3.5 (Protein Metrics Inc.) and the relative amounts of each glycoform at individual sites were determined, as previously described.^31^

### Statistical and survival analyses

All analyses including survival, differential gene expression analysis, and gene set enrichment analyses (GSEA) were carried out and all plots were produced using the R statistical programming language, as described in the **Supplementary methods**. All statistical tests were two-sided. Statistical significance was defined as *P* < 0.05. Kaplan-Meier methods were used to estimate the progression-free survival (PFS; progression/relapse and death of any cause) and the overall survival (OS; death as a result of any cause) in patients that were uniformly treated with R-CHOP-like therapy, as described in the **Supplementary methods**.

## Results

### AGS incidence, distribution and location in DLBCL

The BCCA cohort included 251 and the NCI cohort included 339 primary DLBCL. Their clinical and molecular characteristics, including the full *IGHV-IGHD-IGHJ-IGHC* transcripts are described in the supplemental results (**Tables S1-S5, Figures S1-S4**).

AGS were present in 48% and 55% of GCB-DLBCL (BCCA and NCI cohorts, respectively, **Figure 1A-B**). The majority of the GCB-DLBCL with AGS had at least one in the CDR (84% and 85%). In contrast, only 13 and 18% of the ABC-DLBCL had AGS, and these were CDR+ve for only 7% and 10% of ABC-DLBCL (**Figure 1A-B**).

**Figure 1:**
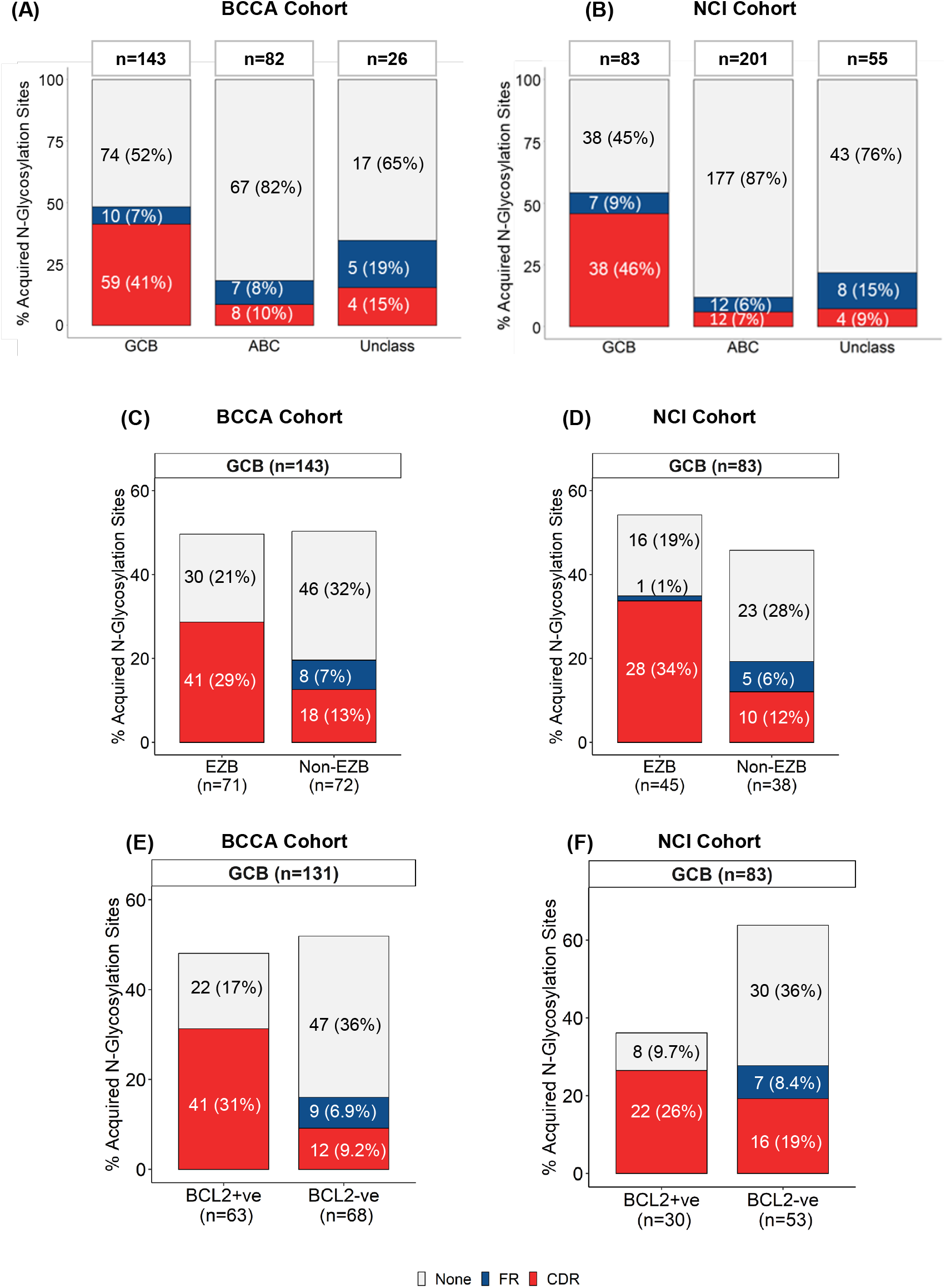
Frequency and location of acquired N-glycosylation sites in the tumor *IGHV-IGHD-IGHJ transcripts of* DLBCL. The tumor *IGHV-IGHD-IGHJ* transcripts of DLBCL were scanned for the presence and location of acquired N-glycosylation sites (AGS). Each case was defined for having at least one AGS in the CDR (red), or AGS in the framework region (FR) only (blue), or no AGS (grey). **(A-B)** AGS frequency and location in DLBCL, divided by cell-of-origin, from the BCCA (**A**) or NCI cohort (**B**). **(C-D)** AGS frequency and location in GCB-DLBCL, divided by Lymphgen EZB subtype or not, from the BCCA (**C**) or NCI cohort (**D**). Also, refer to **Figure S5** for frequencies in the individual non-EZB subtypes. **(E-F)** AGS frequency and location in GCB-DLBCL, divided by cases with a BCL2 translocation (BCL2+ve) or without (BCL2-ve) or not, from the BCCA (**E**) or NCI cohort (**F**).

In the BCCA cohort, 68% of the GCB-DLBCL with AGS in the CDR were EZB, and 32% non-EZB (**Figures 1C, S5A**). However, only 66% of EZB had sites in the CDR while the remaining EZB had acquired no sites. These results were strikingly similar in the NCI cohort, where 73% of the GCB-DLBCL with AGS in the CDR were EZB, and 61% of EZB had CDR-located AGS (**Figures 1D, S5B**).

Prevalence of AGS located in the CDR in GCB-DLBCL with a BCL2 translocation was similar to that in EZB, as expected (65-73%), reflecting 26-31% of all GCB-DLBCL, while the remaining cases with a BCL2 translocation had acquired no sites (**Figure 1E-F**). Conversely, only 31-43% of GCB-DLBCL without a BCL2 translocation had acquired sites, reflecting 16-27% of all GCB-DLBCL, and only 18-30% were located in the CDR (**Figure 1E-F**).

Overall, we identified around 30% of GCB-DLBCL with FL-like features (i.e. with EZB status or BCL2-translocation) and AGS in the CDR, 20% with FL features and no AGS, 20% with no FL features and AGS, and 30% with no FL features and no AGS.

### Mann-type glycans specifically identify low and high-grade lymphomas with FL-like features

Our previous analysis of two lymphoma-derived F(ab)s indicated that Mann-type glycans occupied AGS in the CDR, and not AGS in the framework.^6^ Glycan analysis was performed on 42 F(ab)s from 30 DLBCL and 12 FL with AGS **(Figure 2 and Table S6**). All 12 primary FL cases had AGS in the CDR (**Figure 2A**). The DLBCL samples included 10 with AGS in the CDR and FL features (**Figure 2B**), 20 with AGS in the CDR (12), or in the framework (8) and no FL features (**Figure 2C**).

**Figure 2.**
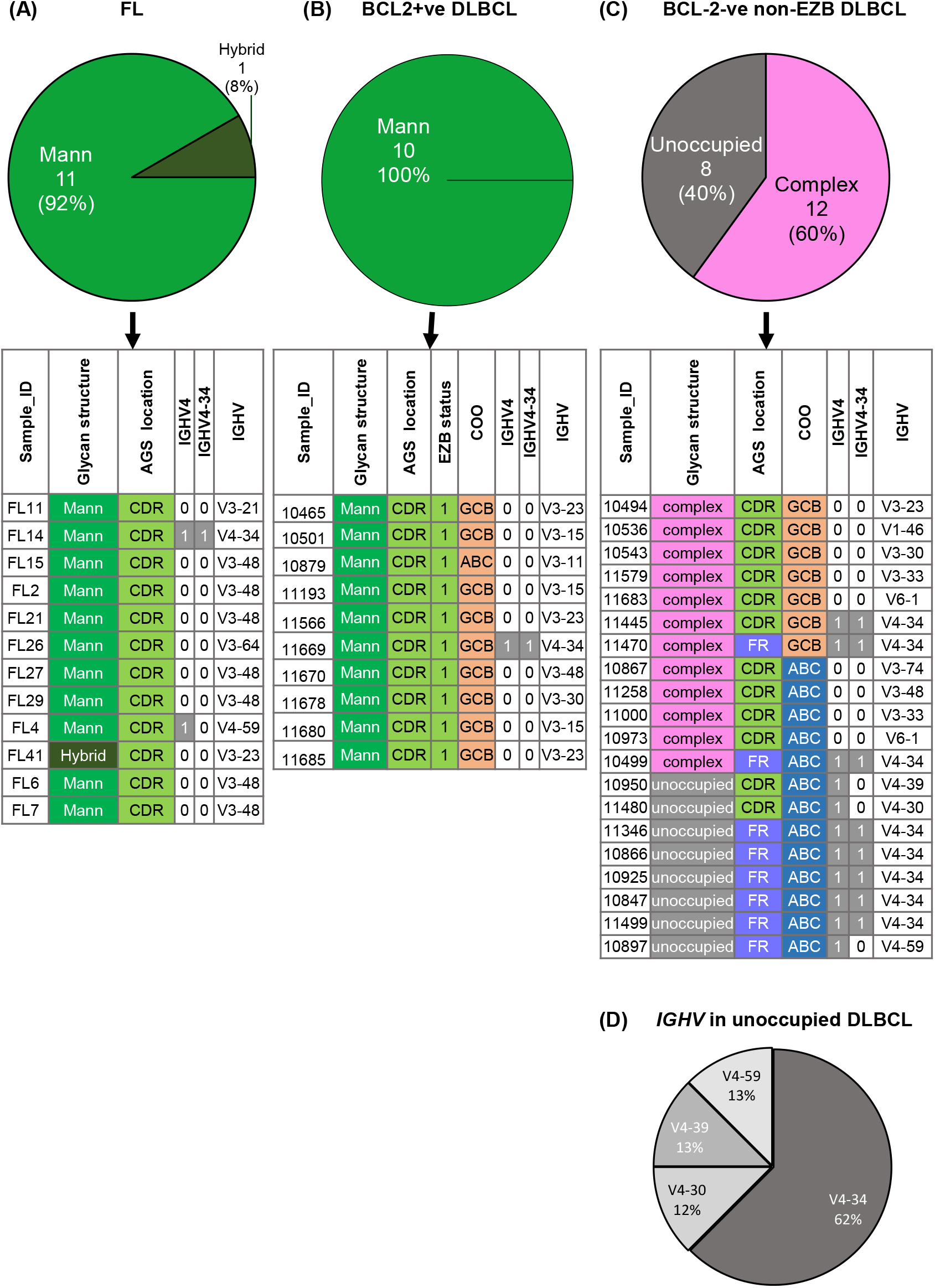
Composition of the glycans occupying the AGS of FL, or DLBCL with and without FL-like genetic features. The glycan composition occupying the AGS in the tumor BCR variable region was determined following the production of recombinant lymphoma-derived F(ab)s and site-specific liquid-chromatography mass spectrometry. Each F(ab) was defined as Mann (oligomannose-type), hybrid, complex, or unoccupied depending on the glycan occupying the AGS in **(A)** FL, **(B)** DLBCL with a BCL2 translocation, or **(C)** without a BCL2 translocation and non-EZB. DLBCL with a BCL2 translocation and an AGS in the CDR were all EZB, GCB-DLBCL and the AGS was occupied by Mann-type glycans. **(D)** *IGHV* use in ABC-DLBCL with the unoccupied AGS. CDR: at least one AGS was located in the complementarity-determining region; FR: AGS were located in framework region only. 0: IGHV4 family or IGHV4-34 gene usage is highlighted as grey (1=yes; 0=no).

Mass spectrometry analysis showed that 10 DLBCL F(ab)s had AGS with Mann-type glycans, 12 had only complex glycans, and 8 had no glycans. All 10 cases with AGS in the CDR and FL features (EZB and BCL2 translocation) were Mann-type F(ab)s (**Figure 2B**). The glycan pattern in these DLBCL resembled FL-derived F(ab)s, also exclusively located in the CDR and Mann-type (**Figure 2A**).

The remaining 20 DLBCL F(ab)s with no FL features had either AGS that were occupied by complex glycans (12) or unoccupied (8) (**Figure 2C**). The 12 complex-type glycans mostly occupied the CDR (10 out of 12), and their glycan composition was irrespective of the cell-of-origin. Instead, the 8 unoccupied AGS were located in the framework (6 out of 8, 75%) and were of F(ab)s derived only from ABC-DLBCL. Interestingly, all the ‘unoccupied’ F(ab)s used *IGHV4* family genes, with 5 out of 8 using *IGHV4-34*. As expected, the natural germine site in IGHV4-34 cases was unoccupied,^6^ but remarkably, the acquired sites in these *IGHV4-34* F(ab)s were also selectively unoccupied (**Figure 2D**).

In summary, only the lymphomas with AGS in the CDR and FL-like features (defined by a BCL2 translocation or EZB status) has Mann-type glycans and shared the features of FL. We called these lymphomas Mann-type DLBCL.

### Mann-type DLBCL identifies lymphomas with short progression-free and overall survival

To understand the clinical significance of glycan types, we investigated progression-free survival (PFS) and overall survival (OS) of Mann-type DLBCL, compared with DLBCL with complex glycans (with an AGS in the CDR and no FL-like features) or unoccupied AGS (with no AGS in the CDR) on the tumor BCR. By analyzing only the cases with BCL2 status available (n=330), we observed that the GCB-DLBCL with AGS in the CDR and a BCL2 translocation had the worse PFS and OS compared to the other DLBCL (**Figure S6**). However, BCL2 translocation is a defining characteristic of the lymphgen EZB subtype,^17,19^ and the use of lymphgen classification for Mann-type DLBCL identification allowed more patients (n=348) for clinical comparisons of Mann-type DLBCL against GCB-DLBCL DLBCL with AGS occupied by complex glycans (non-EZB, with AGS in the CDR) or unoccupied (EZB and non-EZB, without AGS in the CDR) in both the BCCA and NCI cohorts, separately (**Figures S7-8 and 3A-D**). The clinical, genetic and *IGHV* characteristics distribution in these groups are detailed in **Figures S9-S10** and **Tables S7-S8**.

In both cohorts, Mann-type DLBCL had the worst PFS and OS compared to other GCB-DLBCL groups. Remarkably, the PFS and OS of Mann-type DLBCL were similar to those of ABC-DLBCL (**Figures 3A-D, S8**).

**Figure 3.**
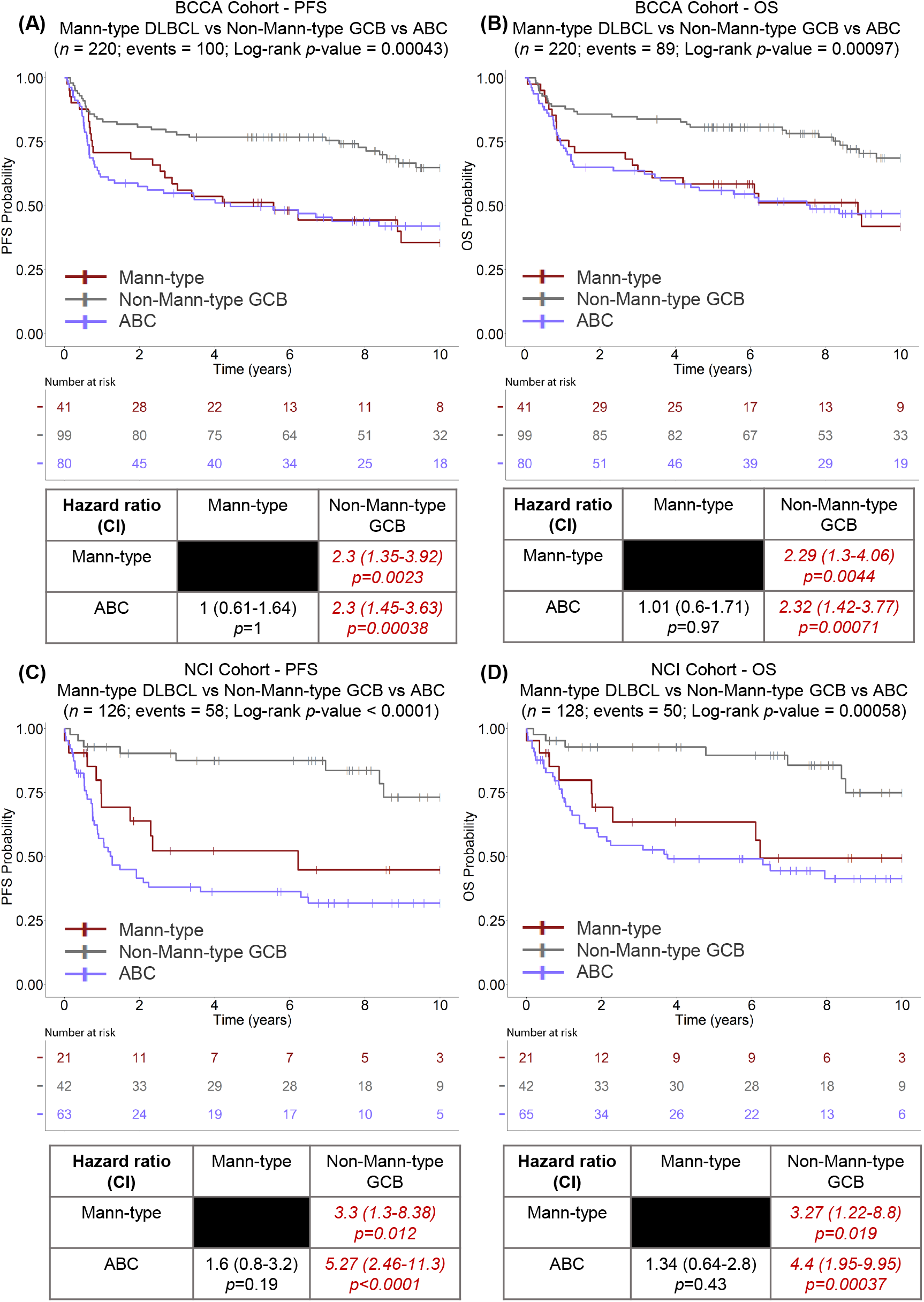
Progression-free survival and overall survival of Mann-type DLBCL. Kaplan-Meyer estimates of progression-free survival (PFS) and overall survival (OS) of Mann-type DLBCL (red) compared with Non-Mann-type GCB-DLBCL (grey) and ABC-DLBCL (purple). **(A)** PFS and **(B)** OS in the BCCA cohort. **(C)** PFS and **(D)** OS in the NCI cohort. Log-rank statistics were used to determine differences between the groups. Hazard ratio values (and 95% confidence interval) and p-values were calculated to compare groups using Cox regression models.

Conversely, Non-Mann-type GCB-DLBCL had favorable outcomes with long PFS and OS, regardless of the occupation or not by complex glycans (**Figure S7-8**).

Since the presence of a *MYC* translocation is known to confer a poor prognosis,^17,23–25^ we used the dark-zone signature (DZsig), a gene signature based on the dual translocation of *BCL2* and *MYC*,^32^ to verify the impact of *MYC* translocation on outcome of Mann-type DLBCL. The worse PFS and OS of Mann-type remained even when comparison to Non-Mann-type DLBCL was done within DZsig+ve or DZsig-ve only (**Figure S11**), indicating an additive impact of Mann-type DLBCL on MYC translocation.

### Mann-type status is an independent prognostic factor of unfavorable outcome

The top five variables significantly associated with worse PFS and OS in univariate analyses (**Figure S12, Tables S9-S10**) were used to assess the independent prognostic value of Mann-type status in multivariate analyses. The covariates included in the analyses were Mann-type status, DZsig, *MYC* rearrangement, *BCL2* and *MYC* dual translocation (HGBL-DH/TH-BCL2), and a high IPI score (**Figure 4**).

**Figure 4.**
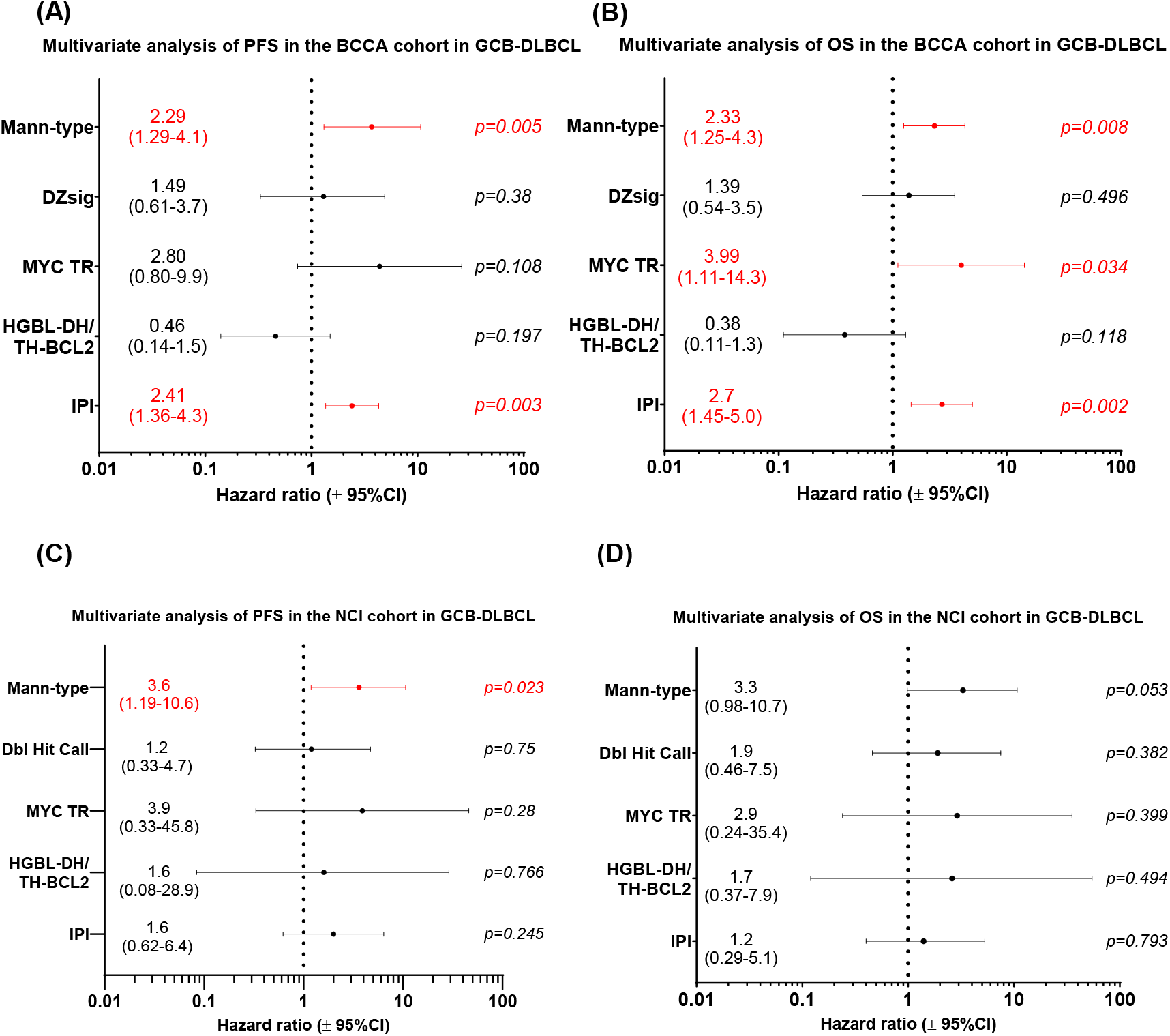
Mann-type status is an independent prognostic factor of PFS and OS in GCB-DLBCL. Multivariate analysis was performed with the clinical and genetic variables that were associated with shorter progression-free survival (PFS) or overall survival (OS) in univariate analyses of GCB-DLBCL. **(A)** PFS and **(B)** OS within GCB-DLBCL of the BCCA cohort. **(C)** PFS and **(D)** OS within GCB-DLBCL of the NCI cohort. The analyses were carried out using Cox regression models and associations were determined using hazard ratios. The variables that independently predicted PFS and OS are in red. Data plotted are hazard ratio ± 95% confidence intervals. Also, refer to **Figure S12** for the univariate analyses of the BCCA and NCI cohorts.

Multivariate analyses in the BCCA cohort revealed that Mann-type status was independently associated with worse PFS and OS (**Figure 4A-B**). Similarly, multivariate analyses in the NCI cohort confirmed that Mann-type status was an independent predictor of shorter PFS and OS (**Figure 4C-D**). A similar multivariate analysis could be performed in the BCCA cohort, but not in the NCI cohort, when Mann-type DLBCL were defined based on AGS in the CDR and BCL2 translocation status only (**Figure S13**). This analysis confirmed the independent prognostic value of Mann-type DLBCL even when BCL2 translocation status only was considered.

### Mann-type DLBCL have enriched MYC, cell cycle, and survival-associated genes

The transcription profile of Mann-type DLBCL was defined by GSEA analyses, using CDR-ve EZB as a comparator (**Table S11**). In the BCCA cohort, 124 genes were significantly upregulated, and 132 genes were significantly downregulated in Mann-type DLBCL **(Figure 5A, Table S11)**. GSEA revealed that Mann-type DLBCL were enriched with MYC signaling, PI3K-Akt-mTORC1 signaling, G2M DNA damage checkpoint, and E2F target-associated genes (**Figure 5B and S14A, Table S12**). Analysis of the NCI cohort validated these results by documenting the enrichment of these gene sets in the Mann-type group (**Table S13**).

**Figure 5.**
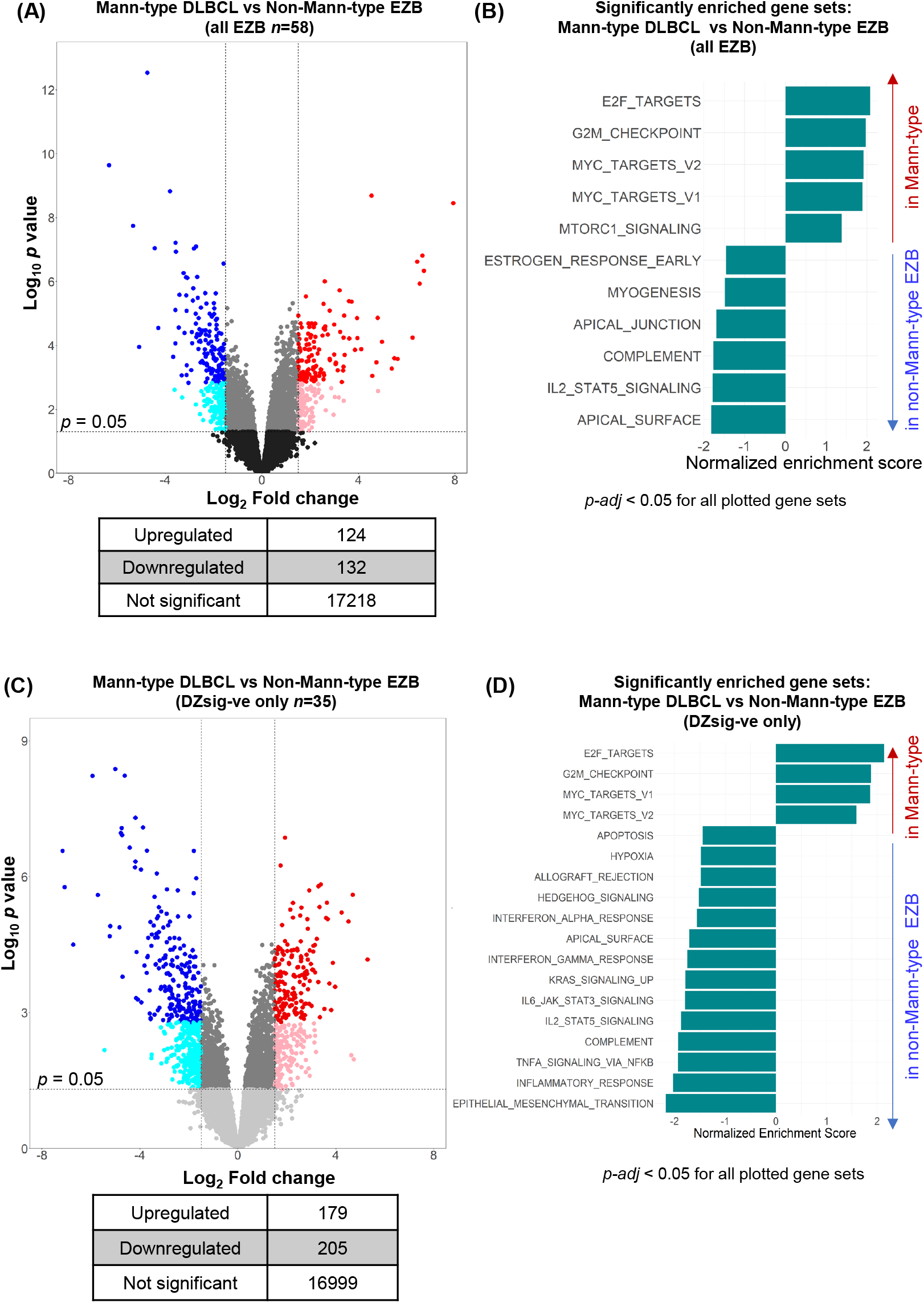
Differential gene expression and gene-set enrichment analysis of Mann-type DLBCL compared to Non-Mann-type EZB. Volcano plots of differential gene expression and GSEA of Mann-type DLBCL compared to Non-Mann-type EZB in the BCCA cohort. **(A-B)** All EZB, **(C-D)** EZB with no dark-zone signature (DZsig-ve) only. **(A, C)** Differential expression was calculated using likelihood ratio tests. Each point in the volcano plots represents a gene with the fold change (FC) and *p*-value for differential expression plotted. A positive log_2_ FC indicates increased expression in Mann-type and a negative log_2_ FC indicates greater expression in Non-Mann-type EZB. Those with a log_2_FC >1.5 or < −1.5 and a corrected *p*-value <0.05 (using Benjamini-Hochberg correction methods for multiple testing) were considered significantly differentially expressed. Those points in light blue and light red had a *p*-value <0.05 and those points in dark blue and dark red had a corrected *p*-value <0.05. The grey points were not significantly different. **(B, D)** GSEA shows the enrichment of Hallmark gene sets in Mann-type or Non-Mann-type EZB. A positive normalized enrichment score (NES) indicates enrichment of the gene set in Mann-type and a negative NES indicates enrichment in Non-Mann-type EZB. Those gene sets that are significantly enriched are represented (Benjamini-Hochberg corrected *p*-value <0.05).

The GSEA analysis was then restricted to EZB with no DZsig to limit the effect of *MYC* translocations on the analysis. Here, 179 genes were upregulated and 205 genes were downregulated in Mann-type DLBCL compared with CDR-ve EZB **(Figure 5C, Table S14)**. GSEA revealed a similar enrichment of MYC signaling, G2M checkpoint, and E2F target genes (**Figures 5D and S14B, Table S15**). These data indicated that the signatures for survival and MYC pathways were associated with Mann-type status, irrespective of the presence of the DZsig signature.

### Mann-type DLBCL have suppressed pro-inflammatory and apoptotic response-associated genes

The GSEA of the downmodulated genes revealed that Mann-type DLBCL had a suppression of multiple gene sets associated with inflammation (**Figure 5, Table S12**). This was most prominent in the lymphomas with no DZsig: the suppressed sets included IFNα, IFNγ, JAK-STAT3, IL2-STAT5, complement, TNFα signaling via NFKB, and inflammatory response (**Figure 5D, Table S15**). Also, analysis of the samples with no DZsig revealed suppression of apoptosis-related genes in Mann-type DLBCL (**Figure 5D, Table S15**). The NCI cohort confirmed suppression of the inflammatory pathways, including complement and IL2-STAT5 signaling-related genes (**Table S13**), despite the small number of cases analyzed in that cohort.

Finally, the mutational profile of Mann-type DLBCL was studied to identify mutations associated with Mann-type DLBCL compared with EZB without AGS in the CDR (**Figure S15**). Overall, no genes were differentially mutated in Mann-type DLBCL compared with CDR-ve EZB.

## Discussion

Our study identifies a distinct type of DLBCL, defined by the occupation of the tumor BCR antigen-binding site by Mann-type glycans. This Mann-type DLBCL type exhibits the shortest PFS and OS compared to other GCB-DLBCL subtypes, which, in contrast, have very favorable outcomes. This finding underscores the need for urgent clinical application of Mann-type DLBCL identification, as it highlights the GCB-DLBCL that do not benefit from conventional immunochemotherapies and establishes Mann-type status as an independent prognostic factor.

Our analysis demonstrates that the simple concomitance of AGS in the CDR and features associated with FL (EZB status or BCL2 translocation) identifies Mann-type lymphomas. Mann-type glycans are not generally found on normal human cells,^33,34^ and are not detected on non-tumor B-cell subsets from blood or lymph nodes.^6^ Therefore, the detection of Mann-type glycans on the sIg antigen-binding site of a clonal B cell population would be highly diagnostic of FL or Mann-type DLBCL. In this study, we utilized a user-friendly pipeline (IgSeqR) to identify the full *IGHV-IGHD-IGHJ* tumor transcript sequence and AGS from RNAseq.^6,29^ Although it is not yet part of routine diagnostics, RNA-Seq offers a cost-effective and accessible alternative to Sanger sequencing, enabling the detection of AGS in the CDR and/or the transcriptional signatures of the EZB genetic subtype in a single procedure.^29^ An alternative diagnostic development could be the detection of DC-SIGN-reactive B cells from flow cytometry of fresh tissue specimens.^6^ In the meantime, the simple combination of immunoglobulin sequencing by Lymphotrack IGH,^35^ and BCL2 translocation tests, which are already in the UK National Genomic Test Directory, would be highly specific for the identification of Mann-type DLBCL and should be implemented in routine diagnostic laboratories.

Our analysis indicates that conventional therapies are unsatisfactory in Mann-type DLBCL and novel strategies are required. While immunochemotherapy combined with BCR-associated kinase inhibitors shows efficacy in ABC-DLBCL subtypes, it does not confer an advantage in GCB-DLBCL subsets,^18^ which include Mann-type DLBCL. Inactivation of N-linked protein glycosylation via oligosaccharyltransferase-B inhibition has been proposed to prevent global N-glycosylation and halt tumor interaction with environmental lectins.^36^ However, this strategy requires optimization to mitigate significant toxicity.

A more feasible and less toxic therapeutic approach involves blocking the Mann:DC-SIGN interaction with an antibody targeting the carbohydrate-recognition domain of DC-SIGN. Anti-DC-SIGN antibodies, developed to modulate immune responses against mannose-bearing pathogens,^37^ breaks pre-existing clusters of Mann-type lymphoma cells with DC-SIGN-expressing cells *in vitro*, depriving the lymphoma cells of a tumor-specific survival signal.^5,6^ However, early intervention at the stage of FL might be even more efficacious. Our studies reveal that FL cells introduce AGS early and glycan’s location and structure remain the same when FL transforms into DLBCL.^6,38^ FL cells have a low proliferation rate and are highly dependent on the interactions driven by environmental DC-SIGN in the tumor lymph node.^5,7^ Here, Mann:DC-SIGN sustains a prolonged low-level pro-survival signal via AKT/PI3K pathway,^8^ which an anti-DC-SIGN antibody can effectively interrupt.^6^

In summary, this study identifies a DLBCL subtype selected by persistent environmental Mann:DC-SIGN interaction, resistant to conventional therapies. We propose novel diagnostic and therapeutic approaches that could significantly impact the early detection and intervention of lymphoma, urging the community to enhance early detection and outcomes for Mann-type DLBCL.

## Acknowledgments

We are very grateful to Freda Stevenson, Professor Emeritus at the University of Southampton, for her continuing mentorship, scientific insight, and support to the group. This work was funded by Cancer Research UK (ECRIN-M3 accelerator award C42023/A29370, and BTERP project C36811/A29101), the Eyles Cancer Immunology PhD scholarship and Fellowship, the Southampton Cancer Research UK Centre and the University of Southampton’s Institute for Life Sciences (M.C. and J.A), the Leukaemia UK John Goldman Fellowship (G.C.) and Pioneer Award (F.F.).

## Authorship

D.T. performed Immunoglobulin Gene analysis, analysed and interpreted data, and wrote the manuscript. M.N. produced lymphoma-derived Fabs, carried out mass spectrometry analysis and contributed to data interpretation. J.A. and J.B carried out mass spectrometry analysis and contributed to data interpretation. B.S. and D.J.B. designed and implemented the IgSeqR Immunoglobulin Gene analysis pipeline, analysed and interpreted data. G.C. analysed and interpreted data. P.D. produced lymphoma-derived Fabs and contributed to data discussion. K.M. contributed to Immunoglobulin Gene analysis of FL samples. K.D. analysed the mutational signatures of Mann+ve DLBCL. D.S. and R.M. analysed and provided the RNAseq data from the BCCA cohort. M.C. designed the glycan analysis studies, supervised research, interpreted data and reviewed the manuscript. F.F. designed the study, supervised research, interpreted data and wrote the manuscript.

