## Supplemental material for "The Origin, Diagnosis, and Prognosis of Oligomannose-Type Diffuse Large B-Cell Lymphoma"

#### List of investigators:

Dylan J. Tatterton*,[1] Maddy L. Newby*,[2] Joel D. Allen*,[2] Benjamin Sale*,[1] Giorgia Chiodin,[1] Patrick J. Duriez,[1] John Butler,[2] Katy J. McCann,[1] David W. Scott,[3,4] Ryan D. Morin,[3] Kostiantyn Dreval,[3] Andrew J. Davies,[1] Dean J. Bryant,[1] Max Crispin^,[2] Francesco Forconi^,[1,5]

[1]School of Cancer Sciences, Faculty of Medicine, University of Southampton, Southampton, United Kingdom;

[2]School of Biological Sciences, University of Southampton, Southampton, United Kingdom;

[3]Centre for Lymphoid Cancer, BC Cancer, Vancouver, BC, Canada;

[4]Department of Medicine, University of British Columbia, Vancouver, BC, Canada;

[5]Haematology Department, Cancer Care Directorate, University Hospital Southampton NHS Trust, Southampton, UK.

*equal contributors as first authors

^equal contributors as senior authors

**Correspondence:** Francesco Forconi, School of Cancer Sciences, Cancer Research UK Centre, Somers Building, MP824, Tremona Road, Southampton, SO16 6YD, UK. **Email:**. **Tel:** +44 (0)23 81205780

#### Supplementary methods

##### Patient cohorts

The BC Cancer Agency (BCCA) DLBCL cohort included cases that met the following criteria: 16 years old or over, treated with curative intent by R-CHOP therapy, with or without radiotherapy, available complete clinical and laboratory, RNAseq data available, and full IG transcript obtained with our pipeline.[1] DLBCL was defined using the 2008 World Health Organization classification as determined by a standardized review by expert hematopathologists.[1] The baseline characteristics and outcomes of the DLBCL in this study were similar to the entire population of DLBCL patients (*n*=1,177) treated with curative intent in BC at that time.[2] The study was reviewed and approved by the University of British Columbia-BCCA Research Ethics Board in accordance with the Declaration of Helsinki, with all patients giving informed consent.

Clinical and laboratory data of the BCCA cohort were obtained from Ennishi et al.[2] and incorporated as individual variables in **Table S1A**. Briefly, the MYC, BCL2, and BCL6 translocations were analyzed by fluorescent in situ hybridization (FISH) on formalin-fixed paraffin-embedded tissue (FFPET) biopsies. Immunohistochemistry staining was performed for MYC, BCL2, CD10, BCL6, MUM1, and ki67, and tumors with ≥30% positive cells were defined as positive. Different cut-offs were used for MYC (≥40%) and BCL2 (≥50%). The COO subtype of the BCCA cohort was determined using the Lymph2Cx 20-gene gene expression profiling assay on the NanoString Platform, which assigned each case to GCB, ABC, or Unclassified DLBCL.[2,3] The LymphGen classification of the BCCA cohort was obtained from Wright et al.[4] To simplify our analysis and minimize the number of LymphGen subgroups with a very small sample size, cases that were called for multiple LymphGen subgroups were grouped as “Comp”. When analysis focused on comparing EZB with non-EZB subgroups, those composite cases that were called for EZB plus at least one other LymphGen subgroup (EZB-Comp) were included in the EZB group. Presence or absence of a BCL2 translocation, as determined by FISH on FFPE biopsies, were obtained Ennishi et al.[2] The dark zone signature (DZsig), a gene signature based on the dual translocation of *BCL2* and *MYC*,[5] and molecular high grade (MHG) classification, were obtained from Davies et al., 2023.[6] The primary sequencing data were downloaded from the European Genome-Phenome Archive (EGA) at https://ega-archive.org/ in the form of aligned BAM files (dataset ID: EGAD00001003783) following approval from the BCCA Data Access Committee.

As a validation cohort, the National Cancer Institute (NCI) cohort of 489 *de novo* DLBCL biopsy samples was used.[7] The NCI cohort was enriched with ABC and Unclassified DLBCL.[7] The biopsies were taken from patients at institutions in the Lymphoma/Leukemia Molecular

Profiling Project (LLMPP) consortium, at the National Cancer Centre of Singapore, or from patients enrolled in the GALGB 50303 clinical trial under institutional review board-approved protocols. The survival data of this cohort was available for the 240 cases treated with R-CHOP or R-CHOP-like therapy. The primary sequencing data were downloaded from the NCI Genomic Data Commons (GDC) portal at https://portal.gdc.cancer.gov in the form of aligned BAM files (accession phs001444.v1.p1). Presence or absence of a BCL2 translocation, as determined by whole exome sequencing, was obtained as summarised tables from the NCI data commons. Other clinical and laboratory data for the NCI DLBCL cohort were obtained from Wright et al.[4] and Schmitz et al.[7] and incorporated into **Table S1B.**

**IG analysis from public RNAseq datasets**

The tumor *IGHV-IGHD-IGHJ-IGHC* rearranged transcripts for both cohorts were generated from bulk RNAseq data using our IgSeqR bioinformatic pipeline.[8,9] The data, downloaded in the form of aligned BAM files, were converted into FASTQ format using SAMtools v1.6. To reduce the computational resources used for *de novo* transcriptome assembly, data filtering was performed by alignment of the FASTQ files to the hg38 reference genome using HISAT2, and only reads that were unmapped or mapped to the *IG* loci as primary or secondary alignments were retained. *De novo* transcriptome assembly was performed using Trinity v2.4.0, with a minimum length of 500 bp and without *in silico* normalization. *IG* loci-derived assembled transcripts were identified using BLASTN against *IGH* reference IMGT/LIGM-DB database and quantified using Kallisto v0.43.1.

**Selection of IG transcripts and identification of acquired N-glycosylation sites**

Following on from the data processing, the *IG* loci-derived assembled transcripts with the highest transcript-per-million (TPM) were aligned to IMGT/LIGM-DB using the IMGT/V-QUEST alignment tool https://www.imgt.org/IMGT_vquest/input. This resulted in an alignment of the unique transcript sequences to germline sequences. Those sequences that did not align with any human *IGH* sequence were discarded. Of the remaining transcripts, one transcript sequence for each sample was selected based on the following criteria, in hierarchical order:

i) The presence of a full *IGHV-IGHD-IGHJ-IGHC* transcript sequence from codon 1 in FR1 to codon 129 in FR4.

ii) 'Productive' V-DOMAIN functionality call. Transcripts with 'no rearrangement found', 'unproductive' and 'no results' calls were discarded.

iii) The most abundant (dominant) *IGHV-IGHD-IGHJ-IGHC* rearranged transcript, defined by having the highest estimated read counts (est. count).

iv) The dominant transcript sequence had to have a 2-fold higher estimated read count than any secondary rearrangements that met the previous criteria.

To avoid excluding unnecessary samples for which a transcript could be selected despite not strictly adhering to all the above criteria, the transcripts were studied on a case-by-case basis to include as many samples as possible. The 2-fold ratio criterion was eased in cases where there was a productive dominant transcript that had a suitable length and high estimated count (>10,000). Using this hierarchical approach, the full *IGHV-IGHD-IGHJ-IGHC* rearranged transcripts were obtained from 251 DLBCL primary samples from the BCCA cohort. The mean est. count was 397,025 ± 16,389 SEM. Of these cases, the survival data was available from 245 cases. The full rearranged transcript sequence was obtained for 339 DLBCL primary samples in the NCI cohort, with a mean est count of 277,829 ± 14,236 SEM.

Acquired N-glycosylation sites (AGS) were identified by manually scanning the deduced amino acid sequence for the N-glycosylation motif (N-X-T/S, where X is any amino acid except proline).[10] The position of the AGS was defined according to the IMGT unique Lefranc numbering system at https://www.imgt.org/IMGTScientificChart/Numbering/IMGT-Kabat_part1.html. The location of the AGS in the CDR or the FR only was defined according to IMGT criteria; those AGS that crossed the CDR and FR border were assigned as being in the CDR.

**IGHV use**

*IGHV* use within DLBCL subgroups was compared to single-cell RNA-sequencing of peripheral blood mononuclear cells from four healthy individuals, two of which were obtained from Bashford-Bashford-Rogers et al.,[11] and the other two from 10X Genomics.[12,13] The means *of the IGHV* frequencies in the four individuals were used as reference values.

**Statistical and Survival analysis**

The statistical analysis of the DLBCL cohorts was carried out using R version 4.2.1 and GraphPad Prism version 8.4.3. Graphical representations of the cohorts were produced using the R package *ggplot2*. Data tables that provide summaries of the DLBCL cohorts were produced using R package *gtsummary.*[14] Frequencies between categorical groups were compared with Pearson's Chi-square test or Fisher's exact tests when appropriate. Continuous variables were compared by T-tests or one-way analysis of variance (ANOVA) followed by post-hoc tests.

Survival analyses were carried out for all patients with survival data available and included patients treated with curative intent at the BCCA using R-CHOP for 3 to 8 cycles, depending on the stage, with or without radiotherapy (BCCA cohort; 245 cases) and at NCI using R-

CHOP or R-CHOP-like therapies (NCI cohort; 155 cases). Progression-free survival (PFS; progression/relapse or death as a result of any cause) and overall survival (OS; death as a result of any cause) were collected as described.[1,7] All endpoints were measured from the point of lymphoma diagnosis. For our analyses, the survival metrics were restricted to 10 years post-diagnosis as the longest time point.

Kaplan-Meier methods were used to estimate PFS and OS, and log-rank tests were performed to compare groups using the R function *survfit* from the *survival* R package. The Kaplan-Meier curves were generated using the *ggsurvplot* R function from the *ggplot2* R package.

Univariable and multivariable Cox proportional hazard regression models were used to evaluate prognostic factors within the BCCA cohort using the R package *survival*.[15] The association of all available genetic and clinical factors with prognosis was analyzed by Cox regression using the *coxph* R function which generated hazard ratios (HRs) for each univariable.

**Multivariate survival analysis**

Those variables that were significantly associated with PFS and OS in univariate analysis ($P<0.05$) were included in the multivariate analyses using the *coxph* function of the *survival* R package, using Cox regression models to calculate the risk, represented using hazard ratios (HRs). [15,16] The covariates included in the analysis had to fulfill the following requirements: (i) for the modeling to be reliable, the number of events per variable (EPV) should not be lower than 10;[17,18] for example, in a subset of data where there were 56 events, no more than 5 covariates should be included in the analysis. (ii) the covariates were statistically associated with survival in the univariate analyses. To not exceed the EPV rule, the covariates were selected based on being significantly associated with both PFS and OS. The 5 most significant variables from the univariate analysis were included in the multivariate analysis. The selection process was made based on the BCCA cohort univariate analysis. The same parameters were included in the analysis with the NCI cohort purely as a validation of the findings in the BCCA cohort, even if the EPV rule was not met due to lower numbers of the GCB-DLBCL population in the NCI cohort. Graphical representation of the multivariate analysis by forest plots was produced using GraphPad PRISM.

**Processing of RNAseq data**

RNAseq data were downloaded in the form of aligned BAM files from the EGA (dataset ID: EGAD00001003783,BCCA cohort), or were obtained via the NCI genomic Data commons for Genotypes and Phenotypes (accession phs001444.v1.p1, NCI cohort). Aligned BAM files were converted to Fastq format using SAMtools *fastq* function.[19] The Fastq files underwent quality control checks using FastQC (Babraham Bioinformatics, Cambridge, UK) and were aligned to the hg38 reference genome using HISAT2 version 2.2.1.[20] Aligned BAM files were checked by FastQC for quality control and were used to assign read counts to genes using HTseq-count against Ensembl GRCh38 v84 gene annotations.[21,22] Overall this pipeline produced a table of gene counts for each sample, which were uploaded into R and merged for further analysis.

##### Differential expression analysis

The tables of gene counts were analyzed for differential gene expression using the R package EdgeR v3.30, following the EdgeR user guide.[23,24] The genes included in the analysis were filtered to remove those with no expression; only genes with > 0.5 counts per million (CPM) in at least 10% of the samples were included in the analysis, and genes with no/low expression were removed. Briefly, the gene expression data were normalized using the Trimmed mean of M-values (TMM). Cox-Reid profile adjusted likelihood ratio (CR) method was used to estimate dispersion, the data were fitted to a generalized linear model (GLM), and differential expression between subgroups was tested using a likelihood ratio test (LRT). These methods resulted in a $\log_2$ fold change (logFC) and a false discovery rate (FDR), which is a corrected P value, for each gene in the analysis. Genes with a log FC ≥ 1.5 or ≤ -1.5 and an FDR < 0.05 were considered significantly differentially expressed.

Gene set enrichment analysis (GSEA) was performed using the fgsea R package from Bioconductor,[25] using the ranked log FC gene list from the differential expression analysis output, and using gene sets from MSigDB v7.2 available from https://www.gsea-msigdb.org/gsea/msigdb.

##### Genetic variant analysis

Published DLBCL genetic variant data for the NCI cohort were downloaded from the National Cancer Institute Genomic Data Commons at https://portal.gdc.cancer.gov/ (accession phs001444.v1.p1) in the form of filtered and annotated MAF files. The BCL2 and MYC translocation data were derived from whole exome sequencing data and downloaded in the form of summarized translocation data.

##### Production of recombinant lymphoma-derived F(ab)s

Recombinant F(ab)s containing the *IGHV-IGHD-IGHJ* and *IGKV-IGKJ/IGLV-IGLJ* sequences from FL and DLBCL cases were produced, with sequences from 35 AGS$^+$ cases of NCI DLBCL cohort and 12 FL from the University of Southampton. [26] The *IG* sequences from the FL and DLBCL tumors were cloned into pDSG plasmids (IBA-LifeSciences, Gottingen, Germany).

The DLBCL *IG* sequences were codon-optimized and the sequences were cloned into the pDSG vectors by Twist Bioscience. The pDSG plasmids encoded the *IGHV-IGHD-IGHJ* with a CH1-hinge and a C-terminal $His_6$ tag, and separate pDSG plasmids encoded the light chain variable region sequences with a *kappa* or *lambda* constant region (**Table S2**). The plasmids encoding the *IGHV-IGHD-IGHJ* or *IGKV-IGKJ/IGLV-IGLJ* rearrangements were co-transfected in a 1:1 molar ratio into MEXi HEK-293E cells (IBA-LifeSciences) according to the IBA LifeSciences protocol, in a 30ml to 200ml scale with a DNA:PEI Max ratio of 1:3. Following incubation of the transfected cells at 37°C for seven days, the supernatants of the transfected cells were collected and were applied to a 250 ml Stericup-HV sterile 0.22 µm filter (Millipore).

The lymphoma F(ab)s were purified from the supernatants on a 5 ml CaptSelect $CH_1$-XL affinity matrix column, on an NGC medium-pressure liquid chromatography system (BioRad, Watford, UK). The elution fractions were pooled and concentrated down to 500 µl volumes using Amicon® Ultra-15 centrifugal filter units (Merck, Darmstadt, Germany).

The F(ab)s were purified further into PBS using a Superdex 200 Increase 10/300 GL gel filtration column (Cytiva). The column was equilibrated with PBS, followed by injection of purified F(ab) material into the NGC chromatography system. The fractions were pooled according to the corresponding peaks on the gel filtration spectra.

#### Supplementary results

##### Characteristics of the cohorts used and *IGHV-IGHD-IGHJ-IGHC* transcripts in DLBCL subtypes

The distribution of cell-of-origin, LymphGen subtypes, clinical or molecular characteristics, and PFS and OS of the BCCA and NCI cohorts used in this study were similar to their extended original cohorts (**Table S3, Figure S1-2).** No skewed *IGHV* use was observed in GCB-DLBCL and EZB (**Table S4-5 and Figure S3**). In contrast, *IGHV4-34* frequency was significantly higher in ABC-DLBCL, and the LymphGen A53, MCD, and “other” subtypes, compared to healthy peripheral B cell cohorts.[11-13] Mean *IGHV* homology to germline was lower in GCB-DLBCL and EZB compared to the other cell-of-origin and lymphGen categories, respectively (**Figure S4**)

In the BCCA cohort, the CDR+ve non-EZB (predicted to be DLBCL with BCR occupied by complex glycans) were 32% of all GCB-DLBCL and included the subtypes ‘other’ (18%), A53 (7%), or ST2 (7%) (**Figure S5A**). In the NCI cohort, the CDR+ve non-EZB were 27% of all GCB-DLBCL ( **Figure S5B**).

### Supplementary figures

#### Figure S1

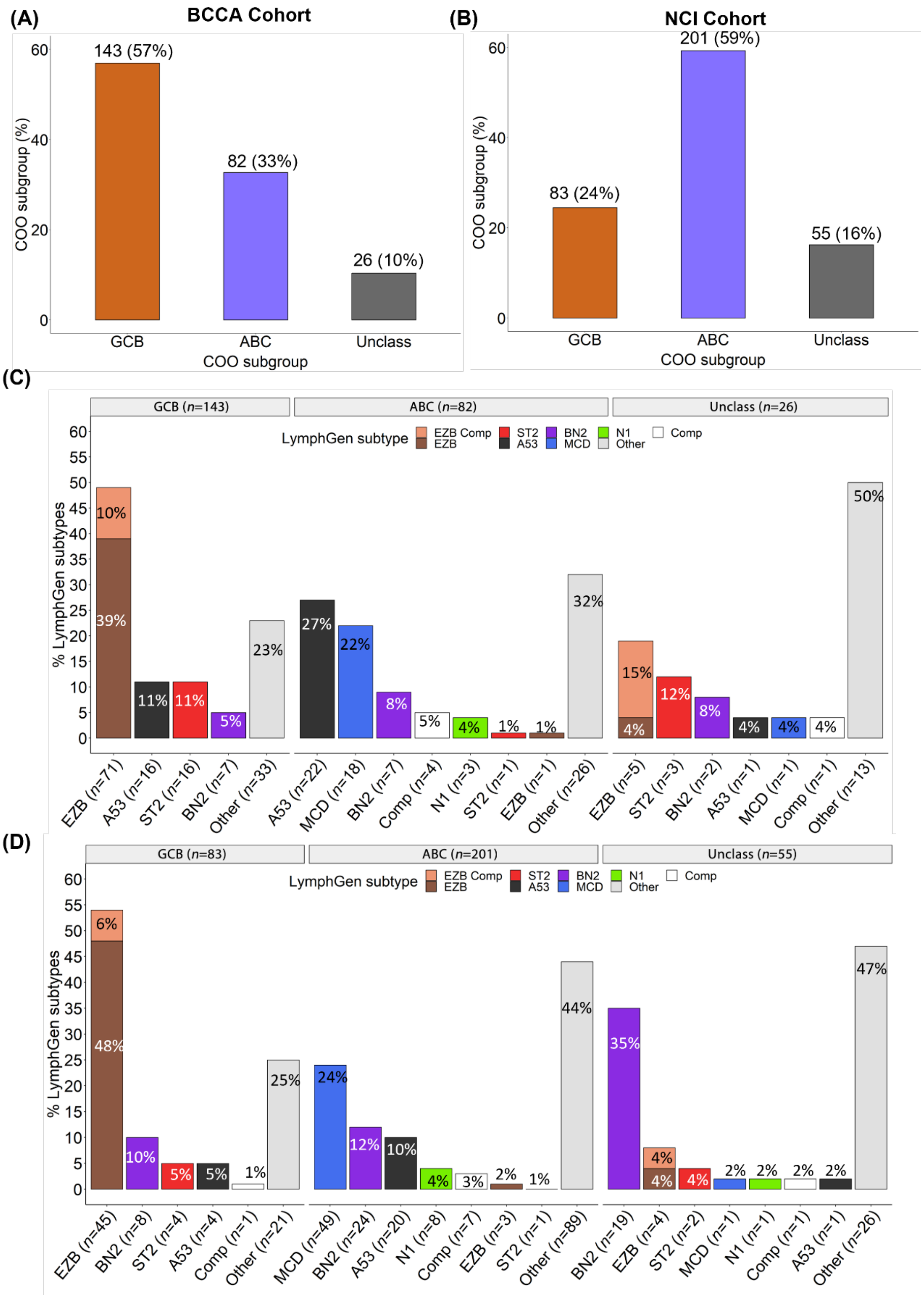


**Figure S1. Genetic and cell of origin characteristics of the DLBCL cohorts used in our study.** The BCCA cohort[2] was used as a discovery cohort and the NCI DLBCL cohort[4,7] as a validation cohort. The full-length tumor *IGHV-IGHD-IGHJ* rearrangements were identified from the DLBCL RNAseq data using our IgSeqR bioinformatic pipeline. Only those DLBCL with the tumor IG rearrangement identified were included in the study. **(A-B)** distribution of the DLBCL in the study as defined by COO in the **(A)** BCCA cohort and **(B)** NCI cohort. **(C-D)** Distribution of the same cases as defined by LymphGenin the **(C)** BCCA cohort and **(D)** NCI cohort.

#### Figure S2

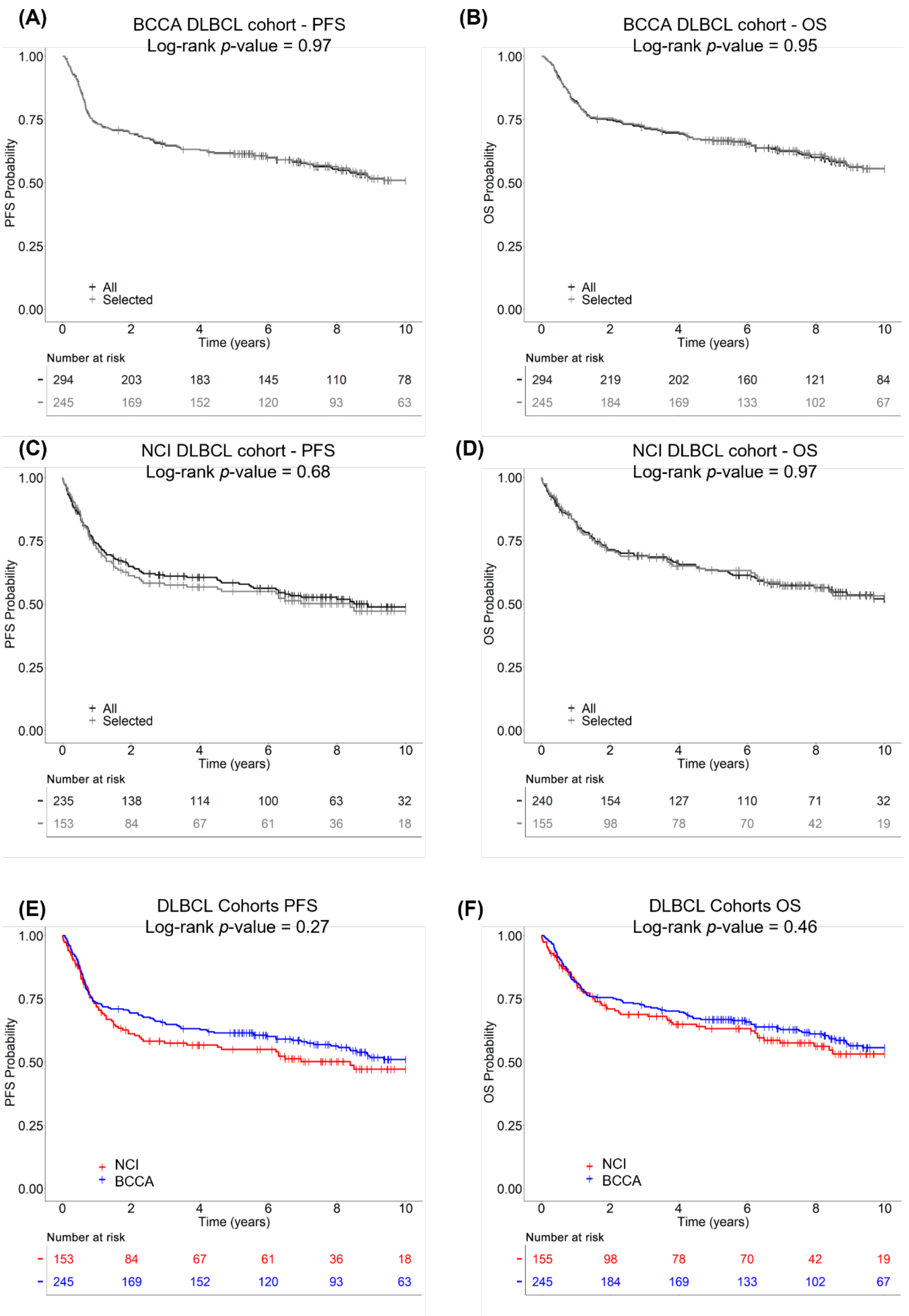


**Figure S2: PFS and OS of the DLBCL selected in the study.** Kaplan-Meier **(A)** progression-free survival (PFS) and **(B)** overall survival (OS) of the patients with *IG* sequences available, used in the present study (selected - light grey), compared with all the patients from the BCCA cohort (all - dark grey). **(C)** PFS and **(D)** OS of the patients with *IG* sequences available, used in the present study (light grey), compared with all the patients from the NCI cohort (dark grey). **(E)** PFS and **(F)** OS of the patients selected in this study from the BCCA (blue) or NCI cohorts (red). The groups were compared by log-rank tests

#### Figure S3

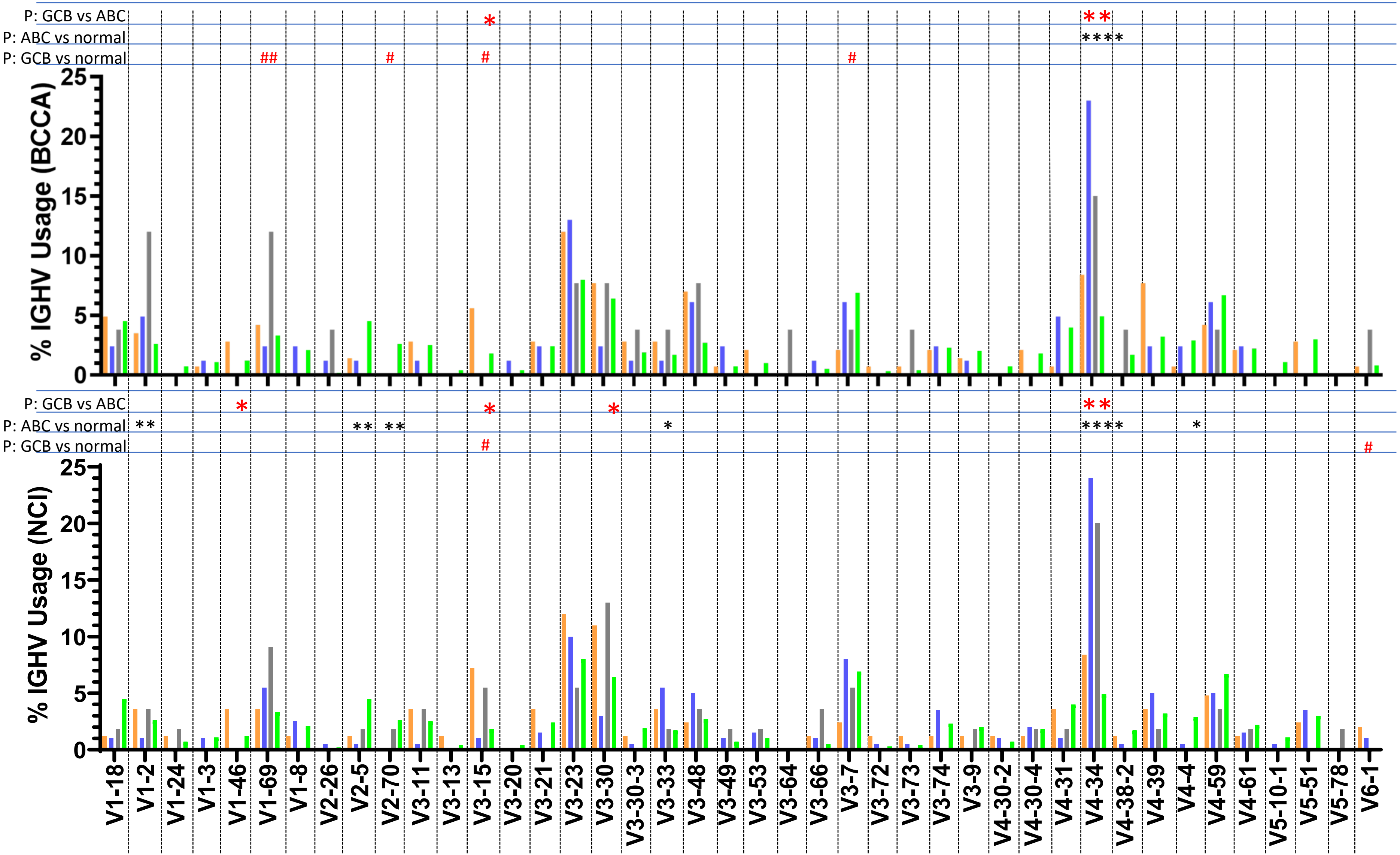


**Figure S3: *IGHV* use in the DLBCL COO subgroups and LymphGen subtypes in the BCCA and NCI cohorts.** Percent *IGHV* usage in GCB (orange), ABC (blue) and Unclass (grey) DLBCL in the BCCA cohort (left panel) and NCI cohort (right panel) compared with IGHV usage in normal peripheral blood mononuclear cells (PBMCs) (green).

#### Figure S4

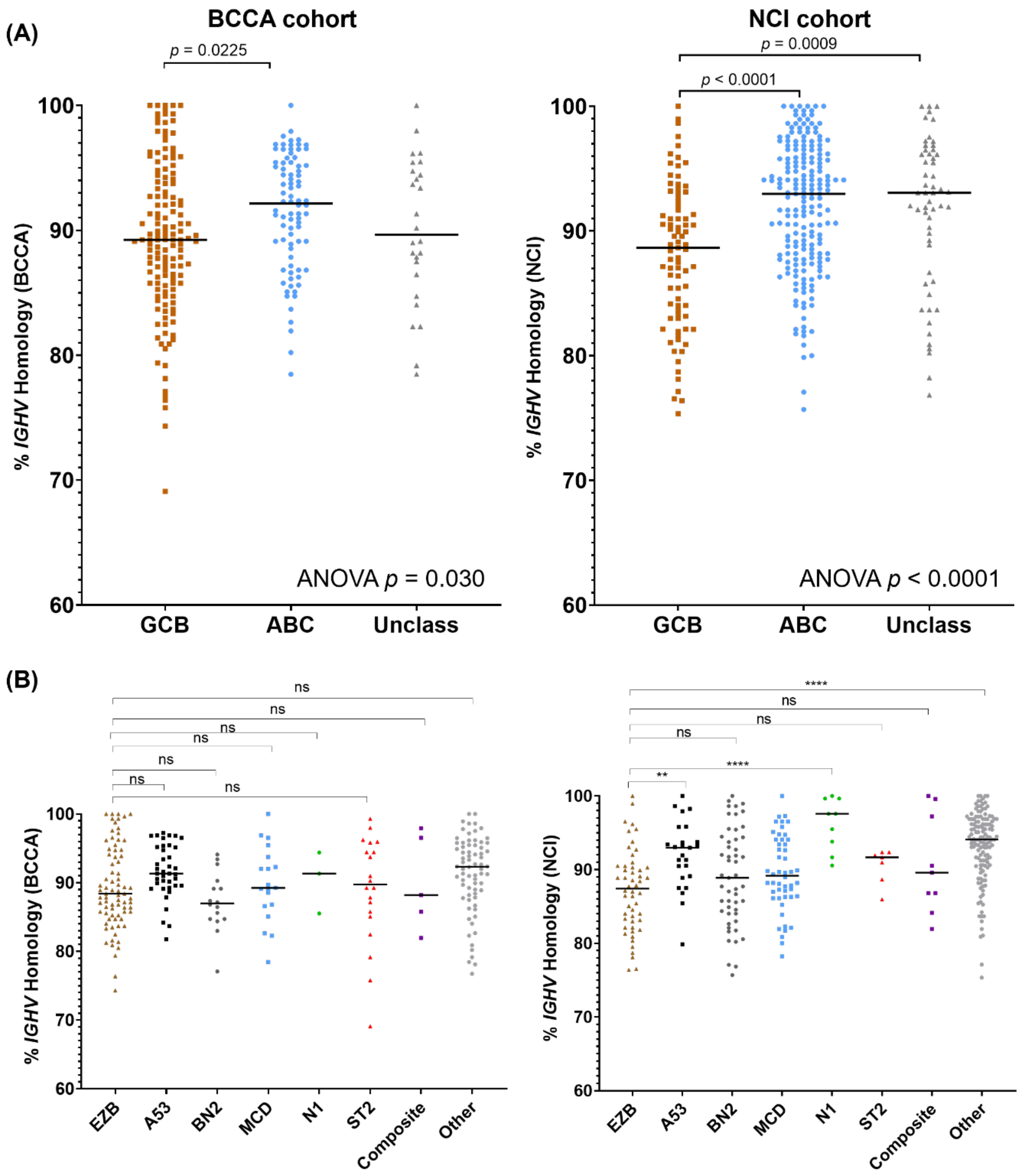


**Figure S4.** ***IGHV* homology to germline in DLBCL. (A)** The percent homology to germline of the tumor *IGHV* sequences in DLBCL from the BCCA cohort (left panel) and NCI cohort (right panel), divided by **COO**. **(B)** The percent homology to germline of the tumor *IGHV* sequences in DLBCL from the BCCA cohort (left panel) and NCI cohort (right panel), divided by **COO**., divided by LymphGen subtype. The *IGHV* homology between the groups were compared by one-way ANOVA and post-Hoc Tukey's multiple comparisons tests. *p<0.05, ** p<0.01, *** p< 0.001, **** p<0.0001.

#### Figure S5

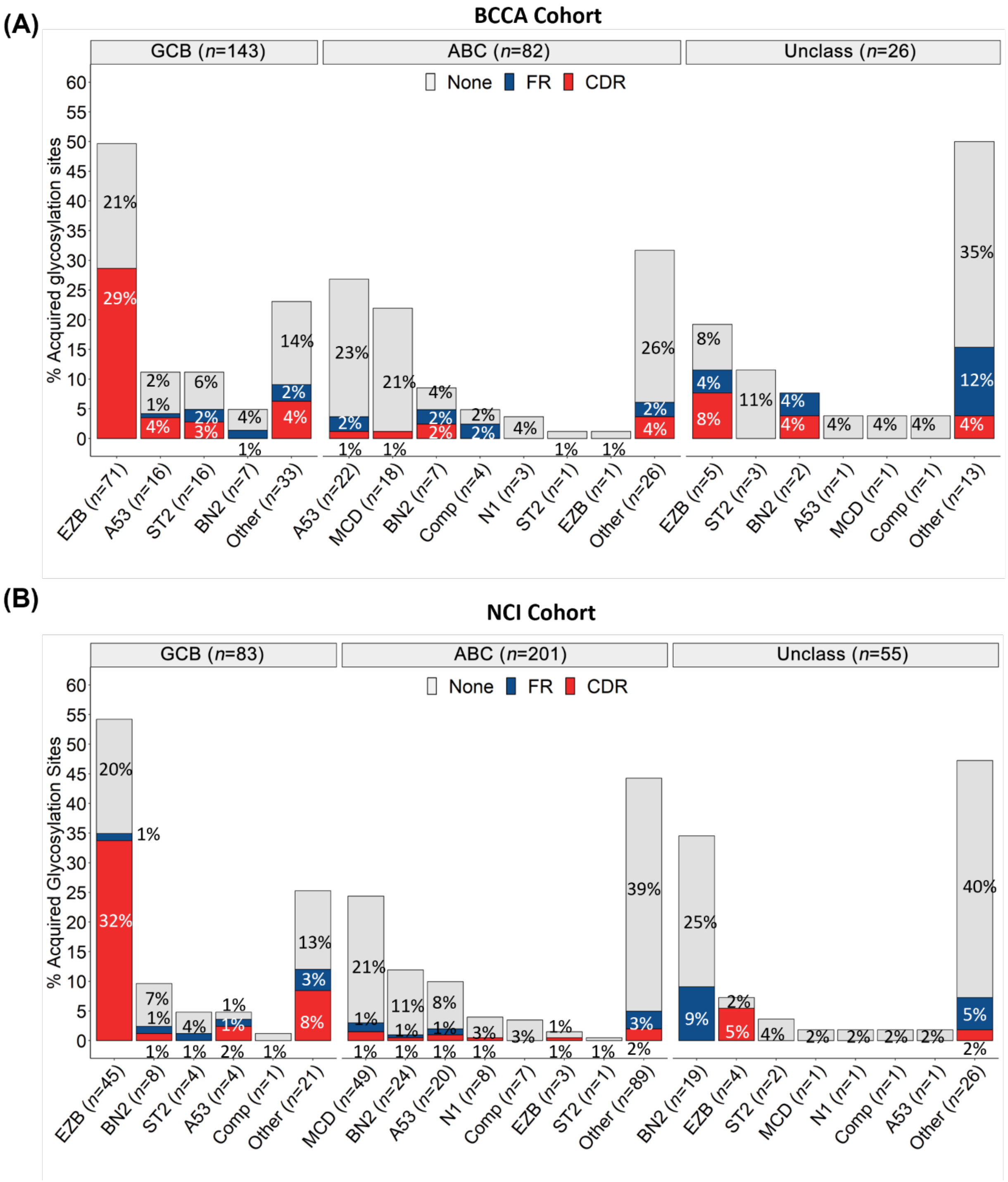


**Figure S5: Frequency and location of the acquired N-glycosylation sites (AGS) in the tumor *IGHV-IGHD-IGHJ* rearrangements of DLBCL divided by of LymphGen subtype.** The tumor *IGHV-IGHD-IGHJ* transcripts were analyzed using the IGMT/V-QUEST numbering system and the derived amino acid sequences were scanned for the presence of acquired N-glycosylation motifs (NxT/S). Each case was defined as having either at least one AGS in the CDR (red), AGS in the FR only (blue), or no AGS (grey). (A) BCCA cohort and (B) NCI cohort.

#### Figure S6

**(A)**

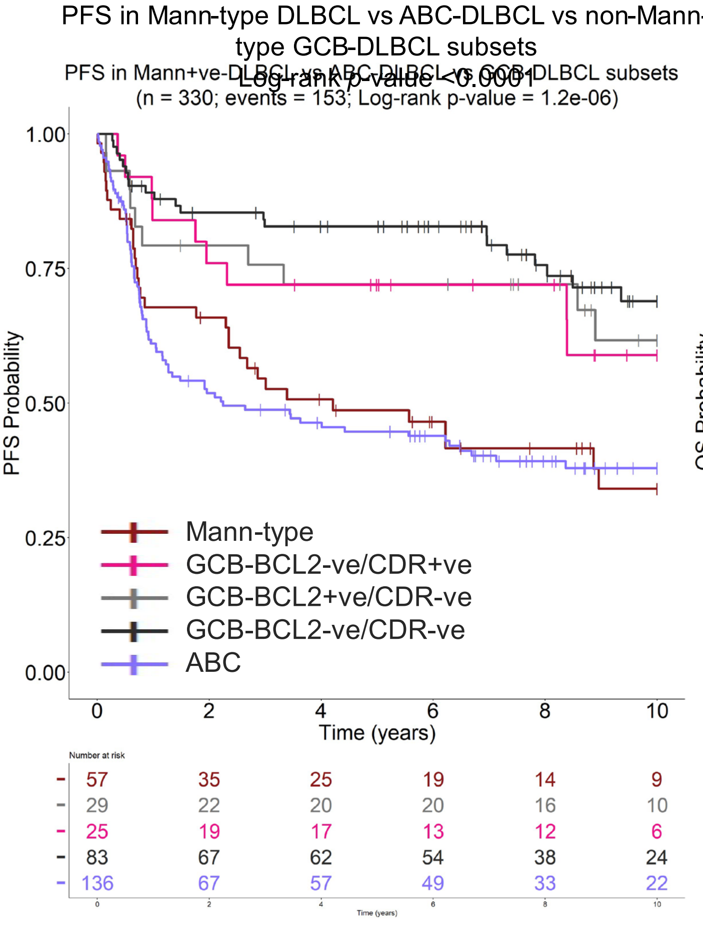


**(B)**

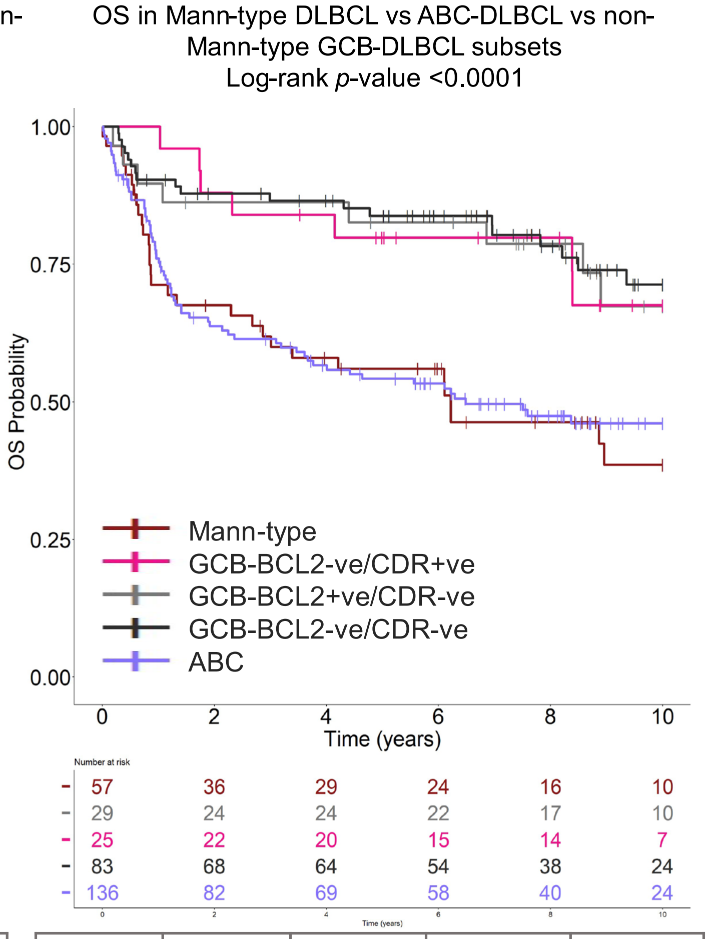


| **Hazard ratio (CI)** | CDR+ve /BCL2-ve | CDR-ve/ BCL2+ve | CDR-ve/ BCL2-ve | ABC |
|---|---|---|---|---|
| Mann-type | 0.48 (0.23-1); *p*=0.05 | *0.45 (0.22-0.92) p = 0.029* | *0.33 (0.19-0.57); p<0.001* | 1.06 (0.71-1.59); *p*=0.78 |
| CDR+ve /BCL2- | | 1.06 (0.43-2.6); *p*=0.9 | 0.68 (0.31-1.49); *p*=0.34) | *2.21 (1.11-4.41); p=0.024* |
| CDR-ve/ BCL2+ve | | | 0.72 (0.34-1.53); *p*=0.4 | *2.34 (1.21-4.52); p=0.011* |
| CDR-ve/ BCL2-ve | | | | *3.24 (2-5.25); p<0.0001* |

| **Hazard ratio (CI)** | CDR+ve /BCL2-ve | CDR-ve/ BCL2+ve | CDR-ve/ BCL2-ve | ABC |
|---|---|---|---|---|
| Mann-type | *0.39 (0.17-0.88); p=0.023* | *0.38 (0.18-0.84); p=0.017* | *0.34 (0.19-0.6); p=0.023* | 0.93 (0.61-1.43); *p*=0.75 |
| CDR+ve /BCL2- | | 1 (0.26-2.76); *p*=1 | 0.87 (0.37-2.07); *p*=0.75 | *2.42 (1.11-5.27); p=0.026* |
| CDR-ve/ BCL2+ve | | | 0.87 (0.38-1.99); *p*=0.74 | *2.43 (1.16-5.05); p=0.018* |
| CDR-ve/ BCL2-ve | | | | *2.78 (1.67-4.63); p<0.001* |

**Figure S9: PFS and OS of Mann-type DLBCL (defined by BCL2 status) is worse than non-Mann-type GCB-DLBCL and similar to ABC DLBCL. (A)** Progression-free survival (PFS) and **(B)** overall survival (OS) of Mann-type DLBCL, DLBCL with complex glycans (CDR+ve BCL2-ve) DLBCL with unoccupied AGS (CDR-ve BCL2+ve, and CDR-ve BCL2-ve GCB-DLBCL), and ABC-DLBCL in the combined NCI and BCCA cohorts. Analysis was performed using Kaplan-Meier methods and a global log-rank statistical test. Cox regression models was used to calculate the hazard ratios.

#### Figure S7

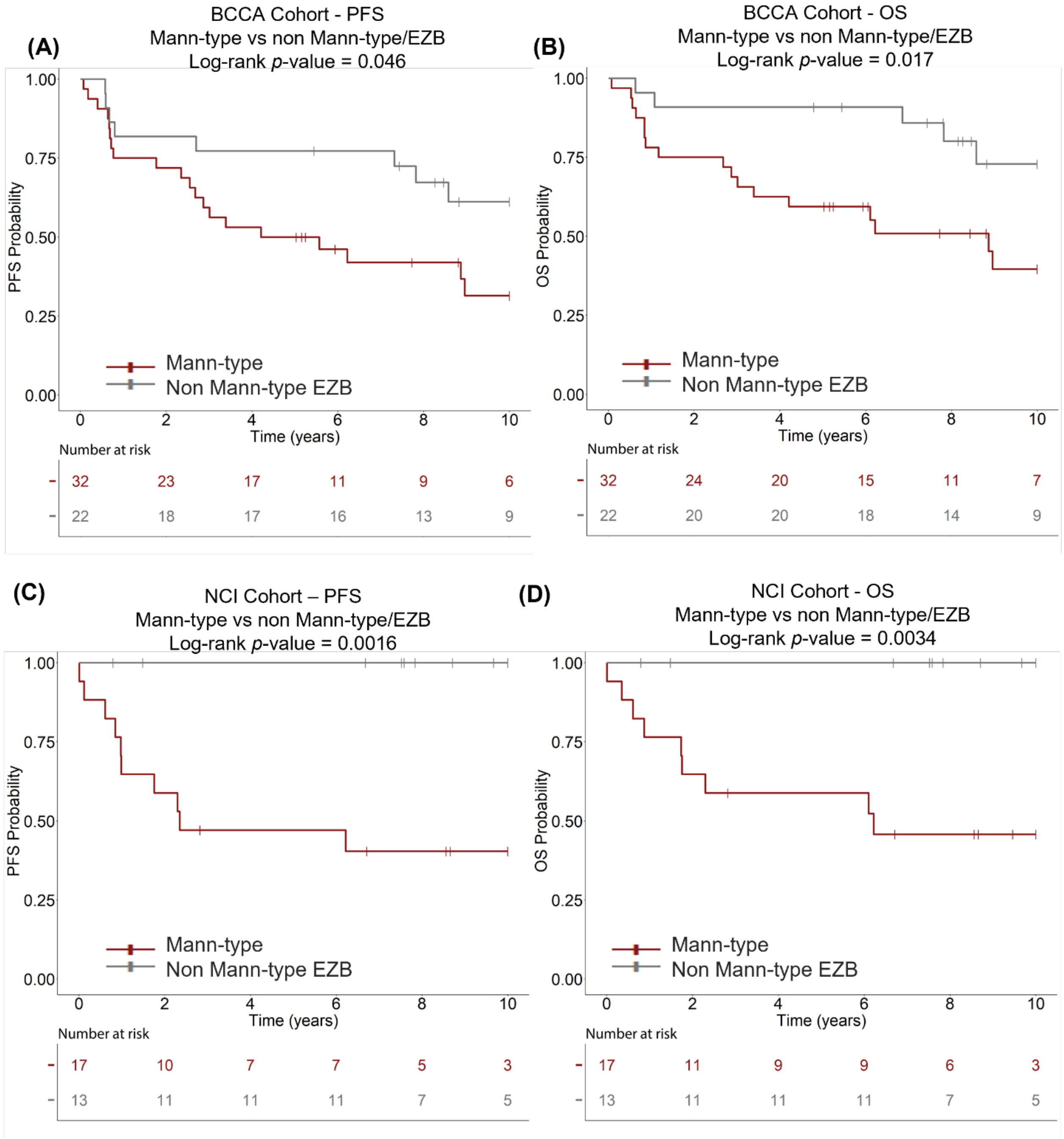


**Figure S7. PFS and OS of Mann-type DLBCL compared to non-Mann-type EZB. (A, C)** Progression-free survival (PFS) and **(B, D)** overall survival (OS) of Mann-type DLBCL (red) compared with CDR-ve EZB (grey), determined by the Kaplan-Meier method, using log-rank statistics for differences between the groups. **(A, B)** BCCA cohort; **(C, D)** NCI cohort.

#### Figure S8

(A) (B)

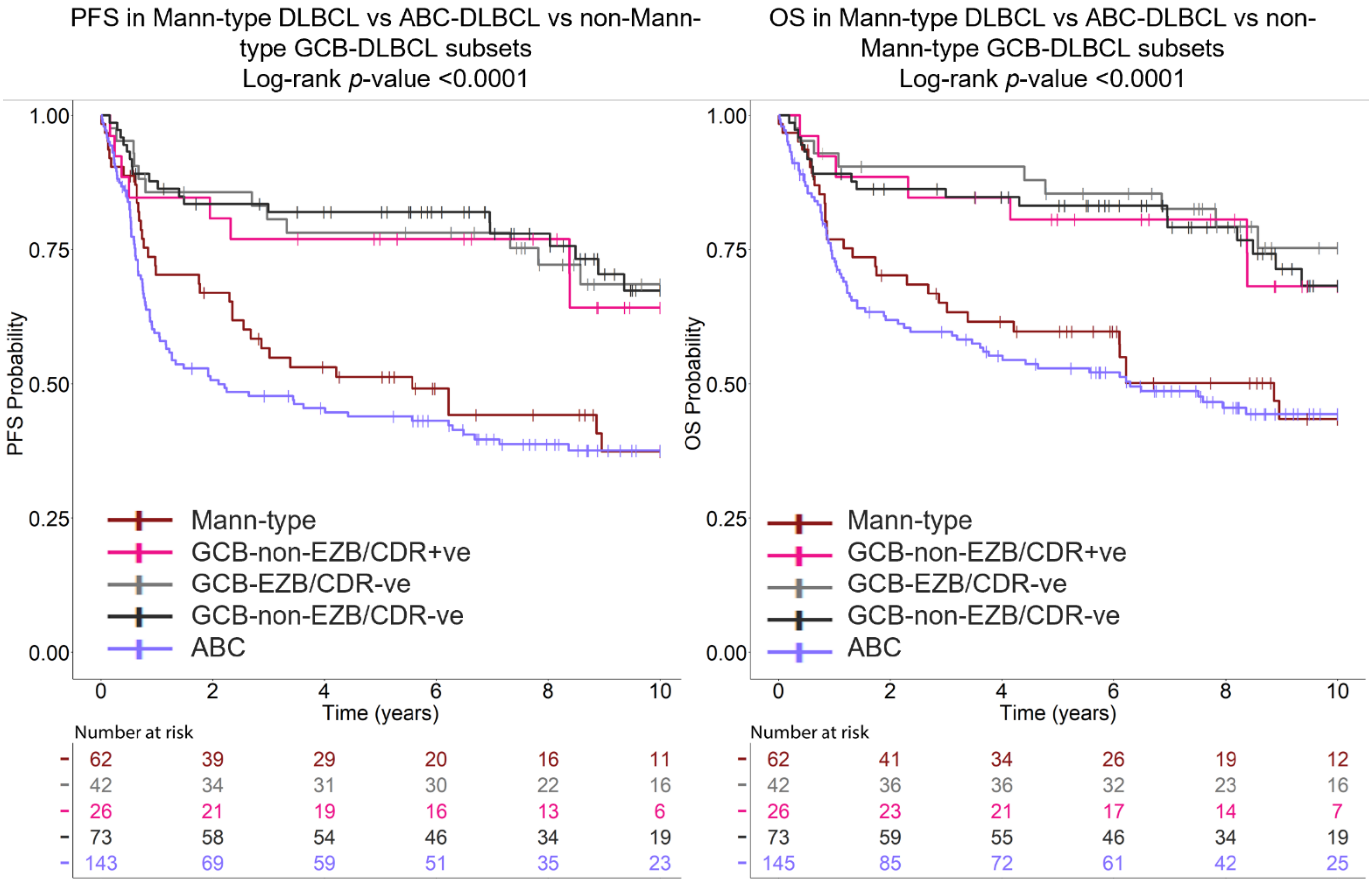


| **Hazard ratio (CI)** | CDR+ve /non-EZB | CDR-ve/ EZB | CDR-ve/ non-EZB | ABC |
|---|---|---|---|---|
| Mann-type | *0.46 (0.21-0.99); p=0.046* | *0.4 (0.21-0.78) p = 0.0068* | *0.38 (0.22-0.66); p=0.0007* | 1.2 (0.81-1.79); *p*=0.37 |
| CDR+ve /non-EZB | | 1.13 (0.46-2.78); *p*=0.78 | 0.83 (0.36-1.89); *p*=0.66) | *2.63 (1.27-5.43); p=0.009* |
| CDR-ve/ EZB | | | 0.94 (0.46-1.94); *p*=0.87 | *2.98 (1.63-5.47); p=0.0004* |
| CDR-ve/ non-EZB | | | | *3.17 (1.93-5.22); p<0.0001* |

| **Hazard ratio (CI)** | CDR+ve /non-EZB | CDR-ve/ EZB | CDR-ve/ non-EZB | ABC |
|---|---|---|---|---|
| Mann-type | *0.44 (0.19-1); p=0.049* | *0.34 (0.16-0.71); p=0.0045* | *0.42 (0.24-0.76); p=0.0038* | 1.13 (0.74-1.72); *p*=0.58 |
| CDR+ve /non-EZB | | 1.29 (0.48-3.47); *p*=0.61 | 0.96 (0.4-2.31);*p*=0.94 | *2.58 (1.19-5.6); p=0.017* |
| CDR-ve/ EZB | | | 1.25 (0.56-2.77); *p*=0.59 | *3.33 (1.66-6.65); p=0.00067* |
| CDR-ve/ non-EZB | | | | *2.67 (1.6-4.47);p=0.00018* |

**Figure S8: PFS and OS of Mann-type DLBCL (defined by EZB status) is worse than other GCB groups and is similar to ABC DLBCL. (A)** Progression-free survival (PFS) and **(B)** overall survival (OS) of Mann-type DLBCL, CDR-ve EZB, CDR+ve non-EZB and CDR-ve non-EZB GCB-DLBCL, and ABC-DLBCL in the combined NCI and BCCA cohorts. Analysis was performed using Kaplan-Meier methods and a global log-rank statistical test. Cox regression models was used to calculate the hazard ratios.

#### Figure S9

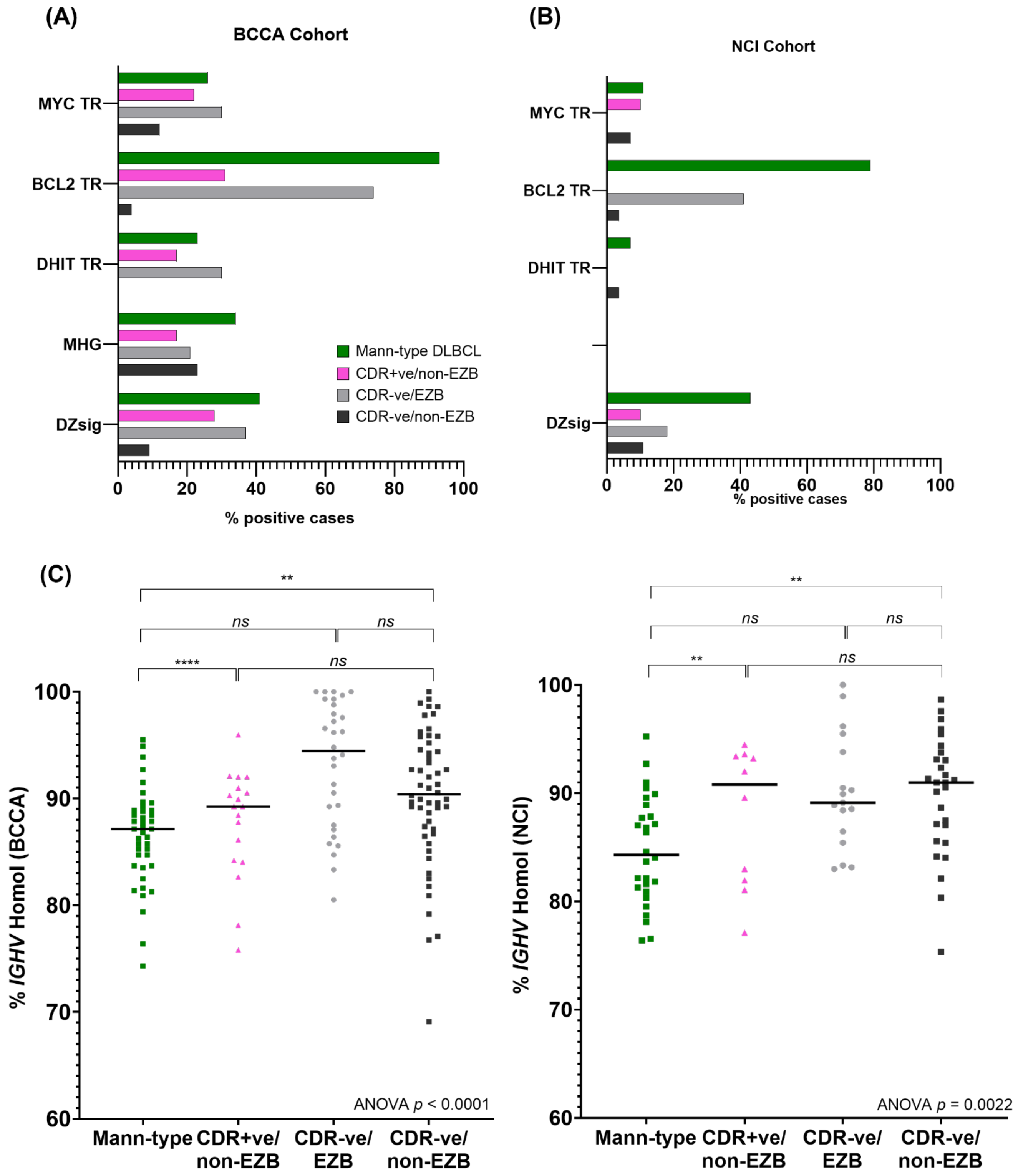


**Figure S9: Genetic and immunogenetic features in Mann-type DLBCL compared with other GCB-DLBCL subsets.** The association of the Mann-type compared to Non Mann-type GCB-DLBCL with MYC translocation (MYC TR), BCL-2 translocation (BCL2 TR) double-hit status (DHIT TR), Molecular High-risk group (MHG), or dark-zone signature (DZsig) in the **(A)** BCCA or **(B)** NCI cohort. **(C)** Tumor *IGHV* percent homology to germline sequence in Mann-type and Non Mann-type (CDR+ve non-EZB, CDR-ve EZB, CDR-ve non-EZB) GCB-DLBCL in the BCCA cohort (left panel) and the NCI cohort (right panel). The homology was compared between groups by one-way ANOVA followed by post-hoc Sidak multiple comparisons test.

**Figure S10**

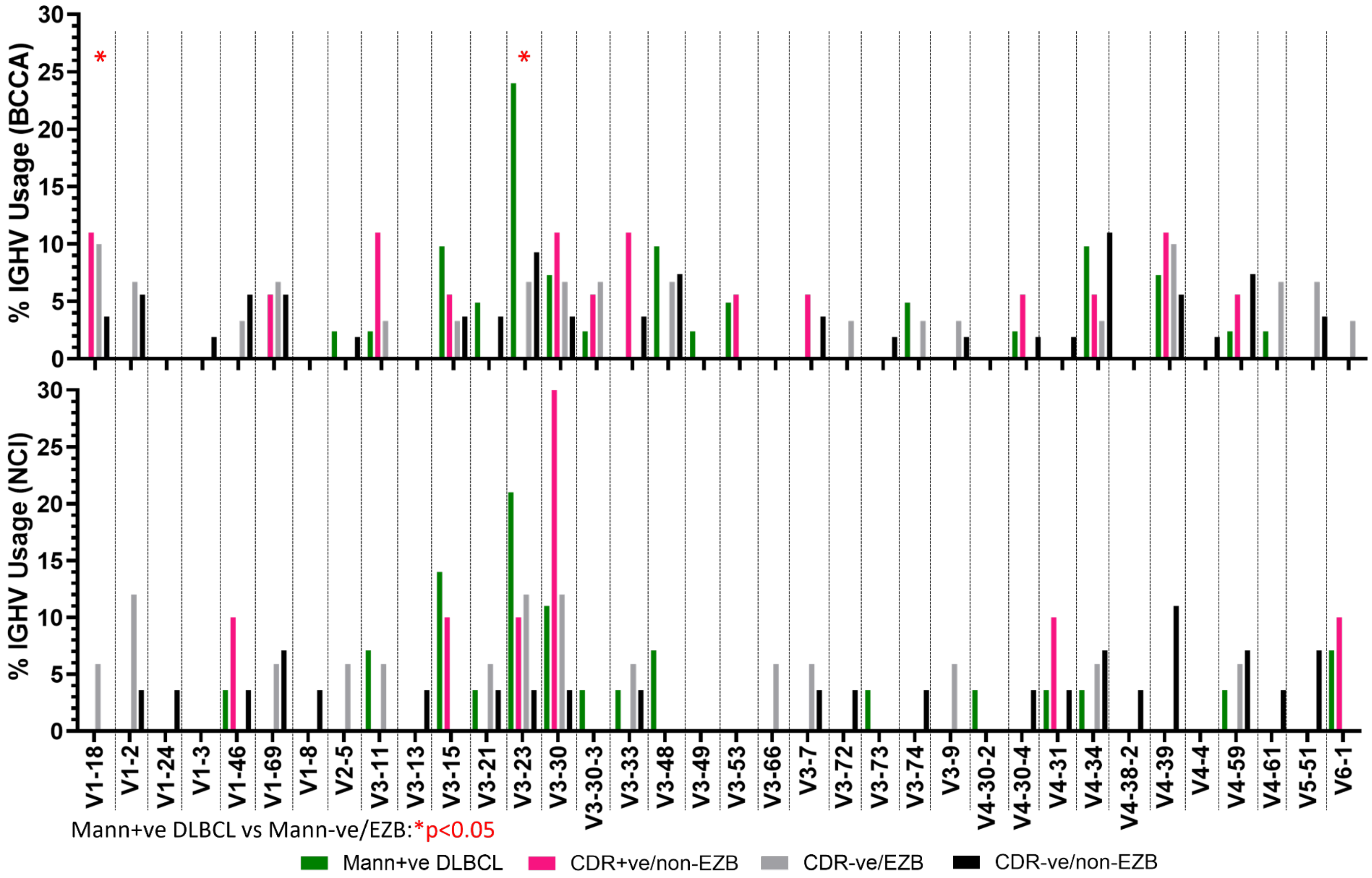


**Figure S10: *IGHV* use in in Mann-type DLBCL compared with other GCB-DLBCL subsets.** Percent *IGHV* usage in Mann-type and Non Mann-type (CDR+ve non-EZB, CDR-ve EZB, CDR-ve non-EZB) GCB-DLBCL in the BCCA cohort (upper panel) and the NCI cohort (lower panel). *IGHV* use frequencies were compared by Fisher's exact test.

#### Figure S11

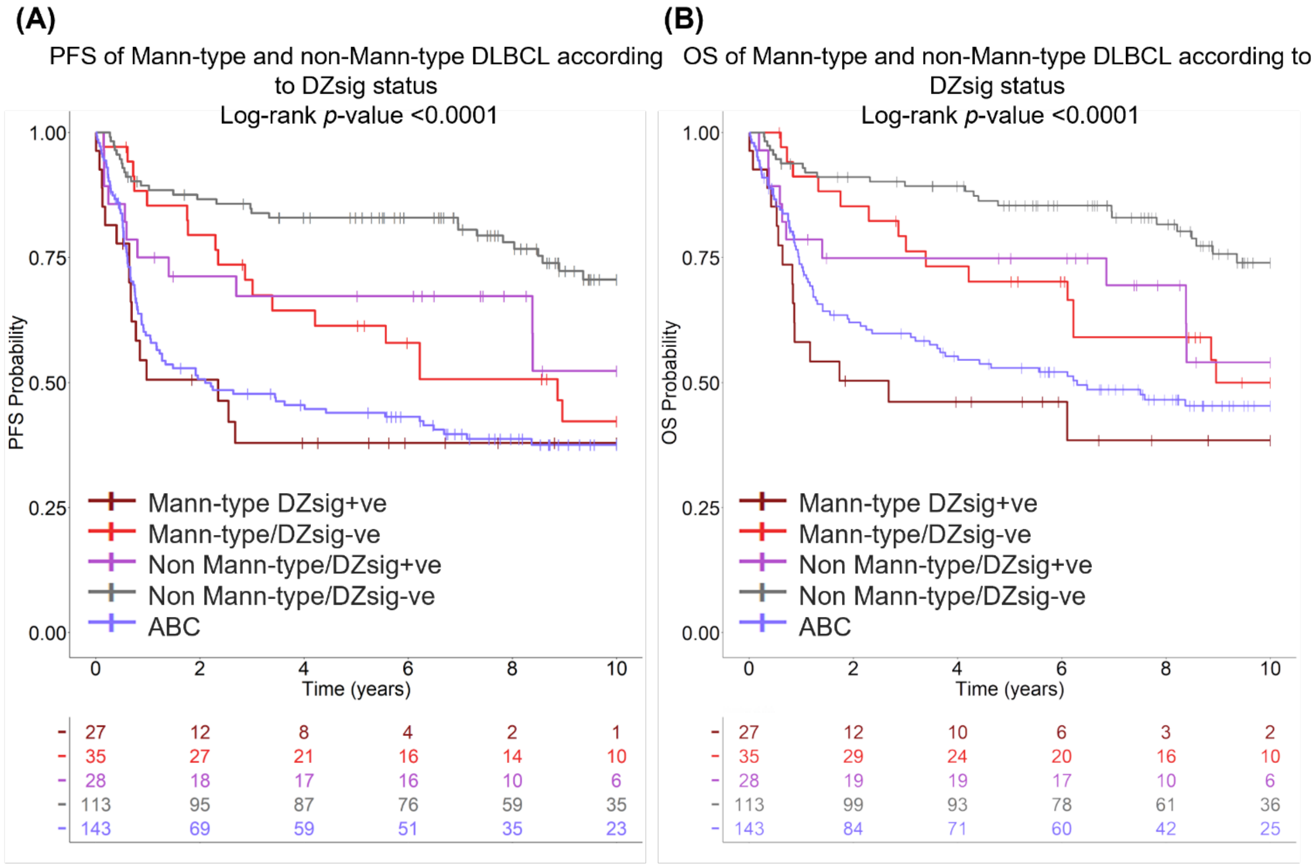


| **Hazard ratio (CI)** | Mann-type /DZ-ve | Non Mann-type /DZ+ve | Non Mann-type /DZ- | ABC |
|---|---|---|---|---|
| Mann-type /DZ+ve | 0.55 (0.28-1.07); p=0.078 | *0.46 (0.21-1); p=0.049* | *0.24 (0.13-0.44); p<0.0001* | 0.83 (0.49-1.42); p=0.51 |
| Mann-type /DZ-ve | | 0.85 (0.4-1.79); p=0.66 | *0.44 (0.24-0.79); p=0.0064* | 1.53 (0.92-2.54); p=0.1 |
| Non Mann-type /DZ+ve | | | 0.52 (0.26-1.04); p=0.065 | 1.18 (0.56-2.5); p=0.66 |
| Non Mann-type /DZ-ve | | | | *0.29 (0.19-0.44); p<0.0001* |

| **Hazard ratio (CI)** | Mann-type /DZ-ve | Non Mann-type /DZ+ve | Non Mann-type /DZ-ve | ABC |
|---|---|---|---|---|
| Mann-type /DZ+ve | *0.48 (0.24-0.99); p=0.046* | *0.45 (0.2-1); p=0.049* | *0.22 (0.12-0.42); p<0.0001* | 0.72 (0.41-1.26); p=0.25 |
| Mann-type /DZ-ve | | 0.93 (0.42-2.07); p=0.85 | *0.46 (0.24-0.87); p=0.018* | 1.5 (0.86-2.62); p=0.15 |
| Non Mann-type /DZ+ve | | | 0.49 (0.24-1.03); p=0.06 | 1.62 (0.83-3.13); p=0.15 |
| Non Mann-type /DZ-ve | | | | *3.28 (2.07-5.21); p<0.0001* |

**Figure S11: PFS and OS of Mann-type and non-Mann-type DLBCL according to DZsig status. (A)** Progression-free survival (PFS) and **(B)** overall survival (OS) of Mann-type/DZ+ve, Mann-type/DZ-ve, Non Mann-type/DZ+ve, Non Mann-type/DZ-ve GCB-DLBCL, and ABC-DLBCL in the combined NCI and BCCA cohorts. Analysis was performed using Kaplan-Meier methods and a global log-rank statistical test. Cox regression models were used to calculate the hazard ratios.

#### Figure S12

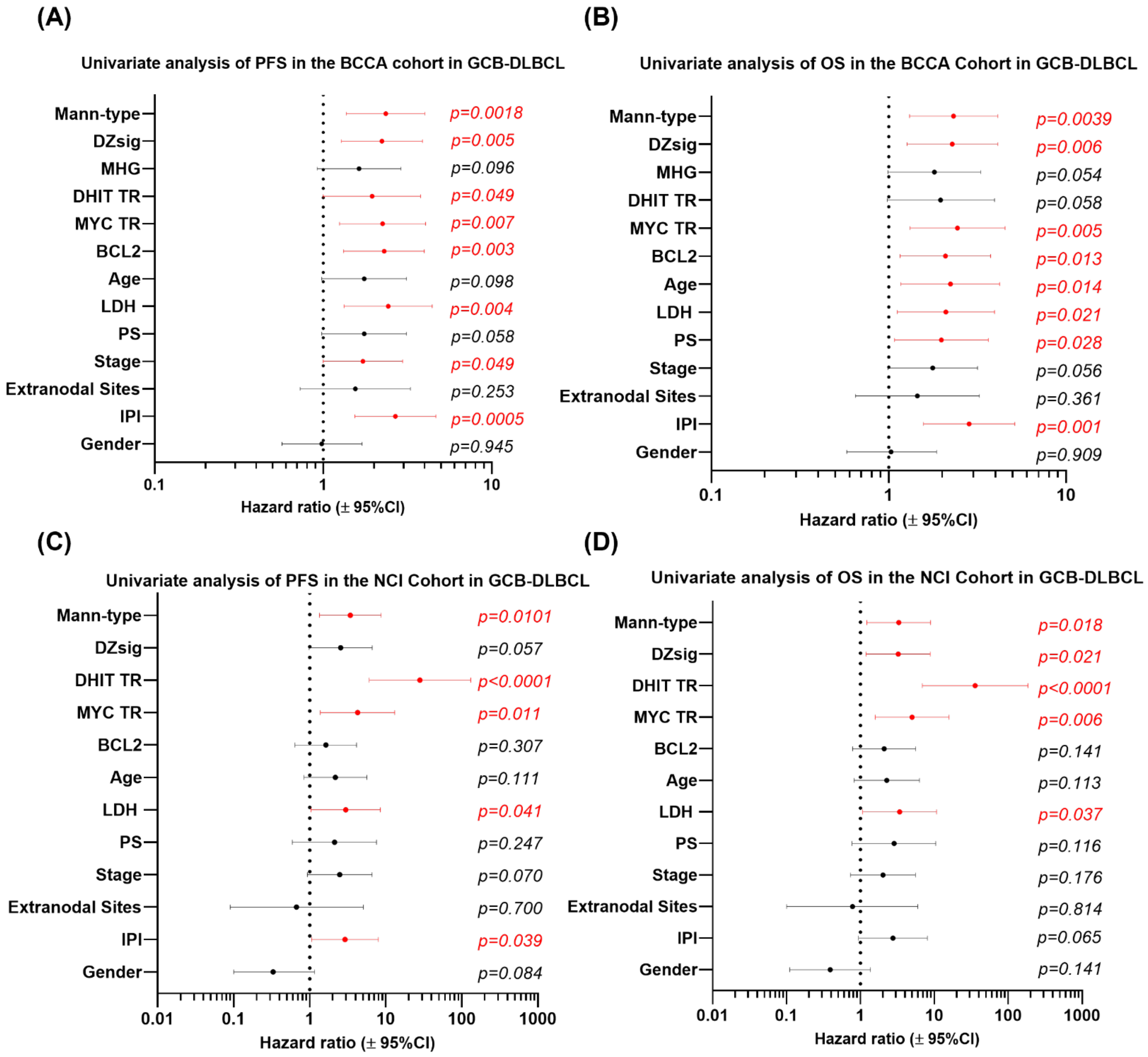


**Figure S12. Univariate survival analysis of PFS and OS in GCB-DLBCL.** The association of Clinical and genetic covariates with (A, C) progression-free survival (PFS) and (B, D) overall survival (OS) in (A, B) the BCCA cohort and (C, D) the NCI cohort in GCB-DLBCL. Data plotted are hazard ratio ± 95% confidence intervals. The analyses were carried out using Cox regression model and associations were determined using hazard ratios.

**Figure S13**

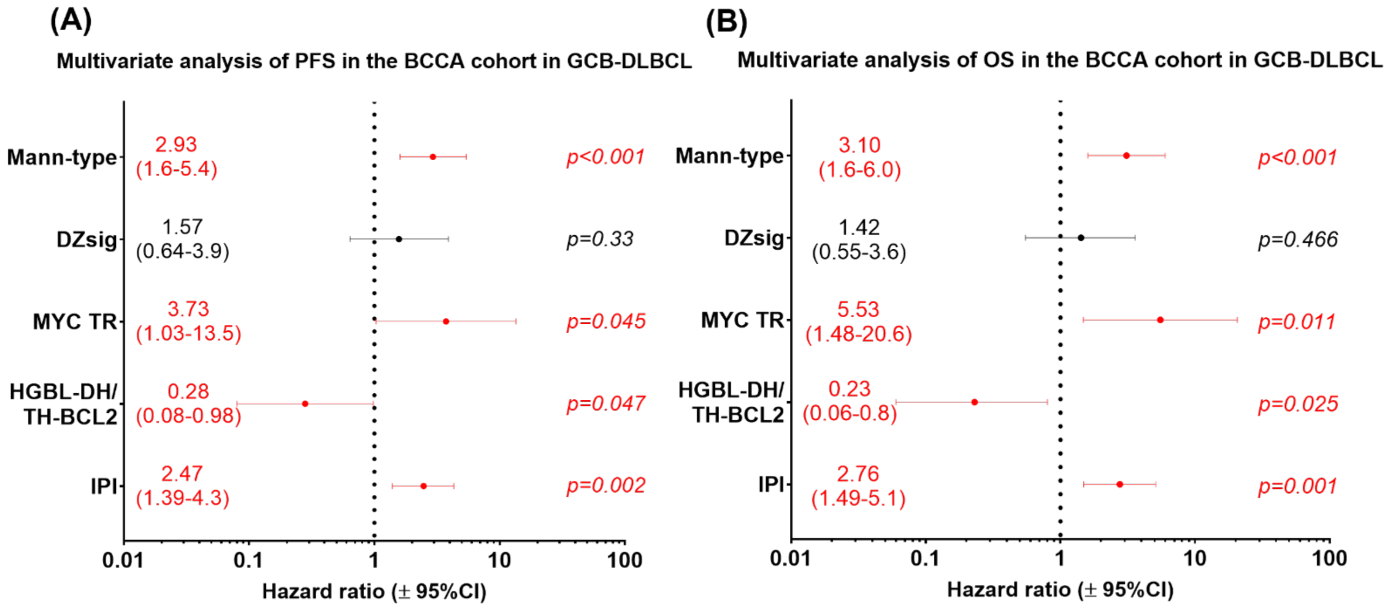


**Figure S13. Multivariate analysis of PFS and OS of GCB-DLBCL. Multivariate analysis was performed with the clinical and genetic variables that were associated with shorter progression-free survival (PFS) or overall survival (OS) in univariate analyses of GCB-DLBCL. (A) PFS and (B) OS within GCB-DLBCL of the BCCA cohort. Mann-type defined as CDR+ve/BCL2+ve The analyses were carried out using Cox regression models and associations were determined using hazard ratios. Data shown are hazard ratio ± 95% confidence intervals. The variables that independently predicted PFS and OS are in red. Also, refer to Figure S12 for the univariate analyses of the BCCA and NCI cohorts.**

#### Figure S14

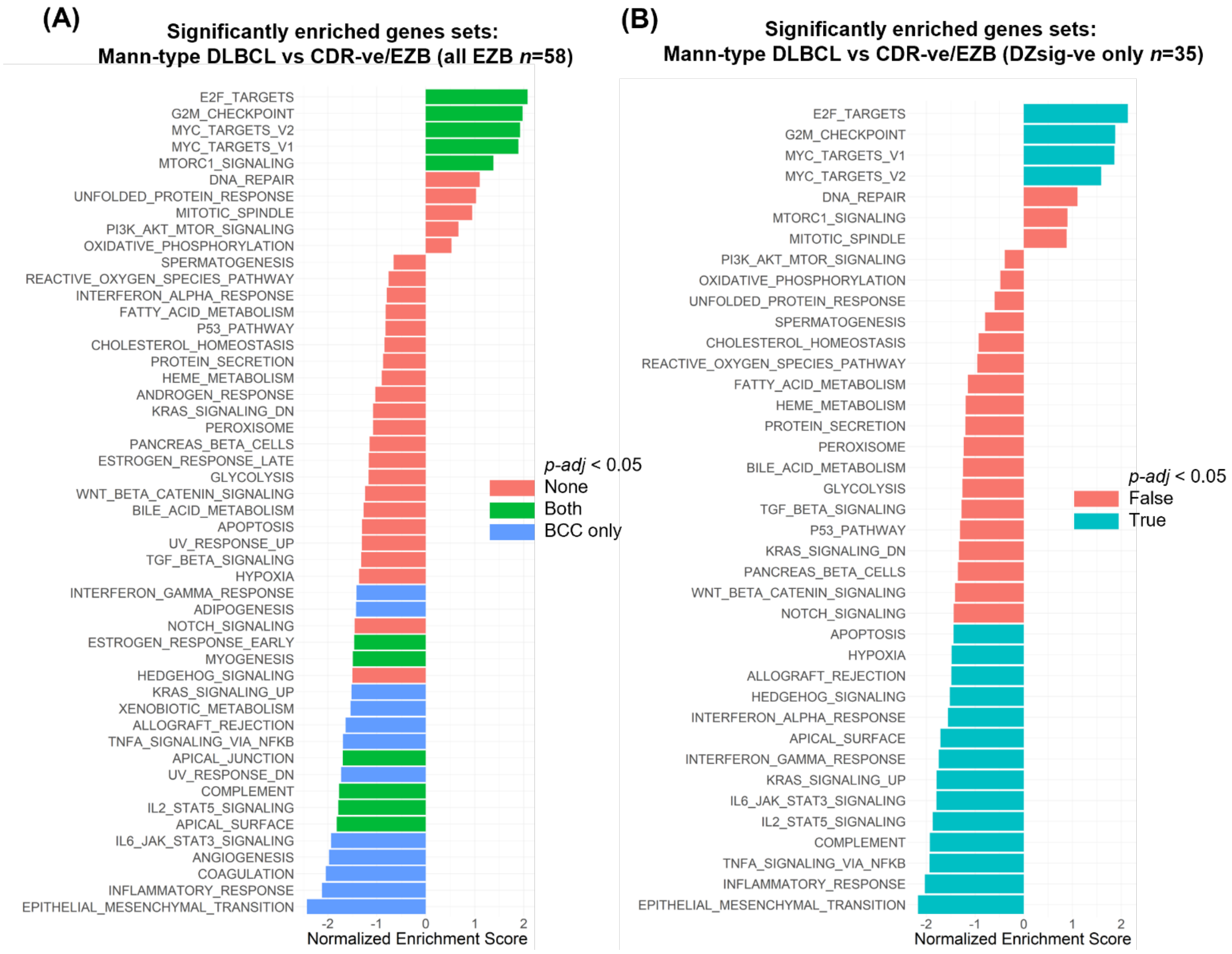


**Figure S14: Gene-set enrichment analysis of Mann-type compared to Non Mann-type EZB in the BCCA cohort.** Gene-set enrichment analysis (GSEA) showing the enrichment of Hallmark gene sets in **(A, B)** Mann-type vs CDR-ve EZB in **(A)** all EZB and **(B)** DZsig-negative EZB. A positive normalized enrichment score (NES) indicates enrichment of the gene set in Mann-type and a negative NES indicates enrichment in CDR-ve EZB. (A) NES plotted for the BCCA cohort. Green bars indicate the gene sets that are significantly enriched in both the BCCA and NCI cohorts (Benjamini-Hochberg corrected p-value < 0.05), blue bars in the BCCA only, red bars not significantly enriched. (B) Analysis conducted in the BCCA cohort; blue bars indicate adjusted $p$<0.05, red bars indicate no significant enrichment

#### Figure S15

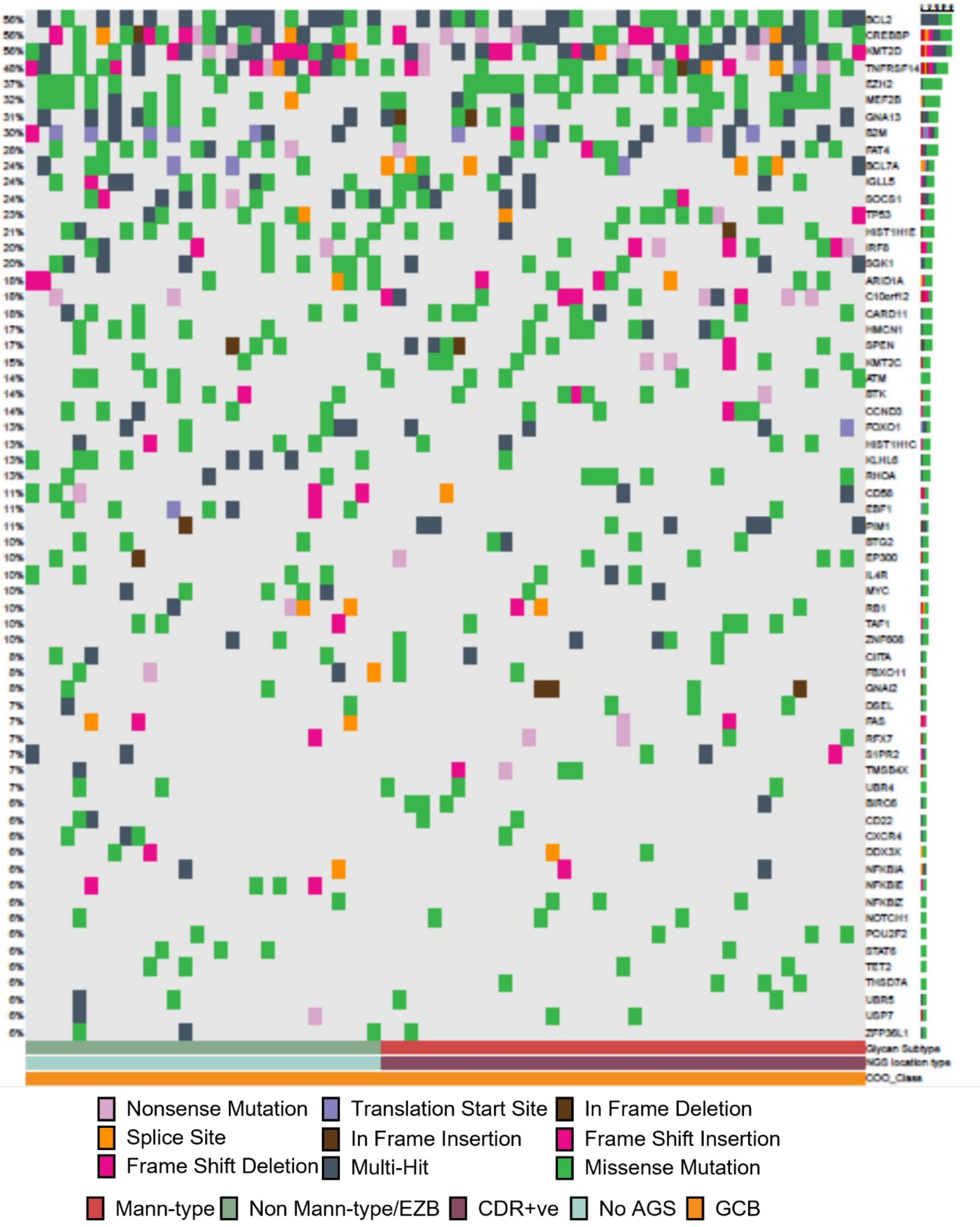


**Figure S15: Oncoplot of simple somatic mutations in DLBCL genes seen in at least 5% of patients with available targeted sequencing data in the BCCA cohort.** Comparison in genetic mutations within EZB lymphomas divided into Mann-type and CDR-ve.

#### Supplementary Tables (Included in a separate Excel file)

##### Table S1A: The clinical, laboratory, genetic, *IGH* characteristics and AGS in the BCCA DLBCL cohort.

Excel spreadsheet showing the genetic and clinical characteristics of the 251 DLBCL from the BCCA cohort in the study. COO: cell of origin (GCB: germinal-center B-cell like; ABC: activated B-cell like; from Ennishi et al.[2]); LymphGen_call: LymphGen classification (Wright et al.[4]). IGHV homol. (%): Percentage homology of the tumor IGHV to germline; AGS: acquired N-glycosylation sites; red characters indicate replacement mutations, generated from RNAseq data the IgSeqR pipeline. DZsig: dark zone signature; MHG: molecular high grade (from Davies et al.[6]); BCL2 TR or MYC TR: presence of BCL2 or MYC translocation; HGBL-DH/TH-BCL2: presence of the *MYC*/*BCL2* double-hit translocations (Ennishi et al.[2]); CPM: Gene expression by counts per million, obtained from gene expression data. PFS; progression-free survival; OS: overall survival (from Ennishi et al.[2]).

##### Table S1B: The clinical, laboratory, genetic, *IGH* characteristics and AGS in the NCI DLBCL cohort.

Excel spreadsheet showing the genetic and clinical characteristics of the 339 DLBCL from the NCI cohort in the study. The characteristics were defined as described in Table S1A. Data were obtained from Schmitz et al.[7] or Wright et al.[4] or were generated from the RNAseq data using the IgSeqR pipeline.

##### Table S2: Sequences inserted into the pDSG vector for the generation of the tumor F(ab) glycoproteins.

Excel spreadsheet showing the sequences inserted into the pDSG vectors for the generation of the tumor F(ab)s. The heavy chain plasmids (pDSG_VH) contained the signal peptide (pHLSeq_SP), the *IGHV-IGHD-IGHJ* sequence, the CH1-hinge sequence, and a 6x His sequence. The light chain plasmids (pDSG_Kappa/pDSG-lambda) contained the signal peptide (pHLSeq_SP), the *IGKV-J//IGLV-J* sequence, and the *IGKC/IGLC* sequence. The restriction enzyme sites and stop codons included in the sequences are also shown in the table.

##### Table S3: Genetic and clinical characteristics of patients from the BCCA and NCI cohorts in the current study.

Excel table showing the distribution of genetic and clinical characteristics in the original cohorts compared to the cohorts in the current study, which included cases where the full *IGHV-IGHD-IGHJ* rearrangements were identified. Data were obtained as described for Table S1A and Table S1B.

##### Table S4: *IGHV* use in COO subgroups of DLBCL compared with healthy peripheral B cells.

Excel spreadsheet showing *IGHV* use in the COO subgroups of DLBCL in the BCCA and NCI cohorts. The *IGHV* use was compared with normal peripheral B cells, obtained from Bashford-Rogers et al.[11] and 10X Genomics.[12,13] COO subgroups were compared with each other and with normal B cells using Fisher's exact tests.

##### Table S5: *IGHV* use in the LymphGen subgroups of DLBCL.

Excel spreadsheet showing *IGHV* use in the LymphGen subgroups of DLBCL in the BCCA and NCI cohorts, compared to normal peripheral B cells by Fisher's exact tests. *IGHV* use was determined as described in Table S4.

**Table S6: Characteristics, genetic lesions and site-specific glycan composition of the DLBCL and FL used for F(ab) studies.**

Excel spreadsheet containing the characteristics, glycan composition, and genetic lesions of the DLBCL and FL tumors used for generation and glycan analysis of the F(abs). The characteristics of the DLBCL cohort were obtained as described in Table S1B. The composition of glycans occupying each AGS was determined using liquid-chromatography mass-spectrometry.

**Table S7: Clinical and genetic characteristics of Mann-type DLBCL.**

Excel table summarising the clinical and genetic features of Mann-type DLBCL, compared with CDR-ve EZB, CDR+ve non-EZB, and CDR-ve non-EZB GCB-DLBCL. The clinical and genetic data of the cohorts were obtained as described in Tables S1A-B. Pearson's Chi-square and Fisher's exact tests were used for statistical analyses.

**Table S8: *IGHV* use in Mann-type DLBCL.**

Excel table summarising *IGHV* use in Mann-type DLBCL, CDR-ve EZB, CDR+ve non-EZB and CDR-ve non-EZB GCB-DLBCL, determined as described in Table S4. *IGHV* use was compared between subgroups using Fisher's exact tests.

**Table S9: Univariate analysis of progression-free survival (PFS) in GCB-DLBCL.**

Excel table showing the association of genetic and clinical parameters with short PFS within GCB-DLBCL in the DLBCL from the BCCA and NCI cohorts. The association of clinical and genetic variables with short PFS in GCB-DLBCL was analyzed using Cox regression models. The parameters included in the analysis were obtained as described in Tables S1A-B.

**Table S10: Univariate analysis of overall survival (OS) in GCB-DLBCL.**

Excel table showing the association of genetic and clinical parameters with short OS within GCB-DLBCL in the BCCA and NCI DLBCL cohorts, analyzed as described in Table S9.

**Table S11: Differentially expressed genes in Mann-type DLBCL in the BCCA cohort.**

Excel table containing the genes differentially expressed in Mann-type DLBCL compared with CDR-ve EZB GCB-DLBCL in the BCCA cohort. logFC: log(fold change); logCPM: log(counts per million); LR: likelihood ratio; FDR: false discovery rate (corrected *p*-value). A positive log Fold Change (FC) indicates increased expression in Mann-type DLBCL and vice versa. Those genes with an FDR<0.05, and a logFc>1.5 or <-1.5 were considered statistically significant differences.

**Table S12: Gene set enrichment analysis (GSEA) of Mann-type DLBCL in the BCCA cohort.**

Table containing the outcome of GSEA with Hallmark gene sets comparing Mann-type DLBCL with CDR-ve EZB GCB-DLBCL in the BCCA cohort. A positive normalized enrichment score (NES) indicates enrichment in Mann-type DLBCL, and vice versa. Those with adjusted *p*-value (*p*-adj)<0.05 were considered significantly enriched. The leading edge shows in order the genes that most drive the enrichment.

**Table S13: Gene set enrichment analysis (GSEA) of Mann-type DLBCL in the NCI cohort.**

Table containing the outcome of GSEA with Hallmark gene sets comparing Mann-type DLBCL with CDR-ve EZB GCB-DLBCL in the NCI cohort, as described in Table S12.

#### Table S14: Differentially expressed genes in Mann-type (DZsig-negative cases only) in the BCCA cohort.

Table containing genes differentially expressed in Mann-type DLBCL vs CDR-ve EZB (within DZsig-negative GCB-DLBCL) in the BCCA cohort. Data are as described in Table S11.

#### Table S15: Gene set enrichment analysis (GSEA) of Mann-type DLBCL in the BCCA cohort, restricted to DZsig-negative cases.

Table containing the outcome of GSEA with Hallmark gene sets comparing Mann-type DLBCL with CDR-ve/EZB cases in the BCCA cohort, restricting the analysis to DZsig-negative cases of GCB-DLBCL. Data are as described in Table S12.
